# The Up state of the SARS-COV-2 Spike homotrimer favors an increased virulence for new variants

**DOI:** 10.1101/2021.04.05.438465

**Authors:** Carolina Corrêa Giron, Aatto Laaksonen, Fernando Luís Barroso da Silva

**Affiliations:** Universidade de São Paulo, Departamento de Ciências Biomoleculares, Faculdade de Ciências Farmacêuticas de Ribeirão Preto, Av. café, s/no – campus da USP, BR-14040-903 Ribeirão Preto SP, Brazil; Universidade Federal do Triângulo Mineiro, Hospital de Clínicas, Av. Getúlio Guaritá, 38025-440 Uberaba, MG, Brazil; Department of Materials and Environmental Chemistry, Arrhenius Laboratory, Stockholm University, SE-106 91 Stockholm, Sweden; State Key Laboratory of Materials-Oriented and Chemical Engineering, Nanjing Tech University, Nanjing, 210009, PR China; Centre of Advanced Research in Bionanoconjugates and Biopolymers, Petru Poni Institute of Macromolecular Chemistry, Aleea Grigore Ghica-Voda, 41A, 700487 Iasi, Romania; Department of Engineering Sciences and Mathematics, Division of Energy Science, Luleå University of Technology, SE-97187 Luleå, Sweden; Department of Chemical and Biomolecular Engineering, North Carolina State University, Raleigh, NC 27695, United States

**Keywords:** SARS-CoV-2, mutations, conformational states, coronavirus, electrostatic interactions, epitopes, binding affinity, protein-protein interactions

## Abstract

The COVID-19 pandemic has spread widely worldwide. However, as soon as the vaccines were released – the only scientifically verified and efficient therapeutic option thus far – a few mutations combined into variants of SARS-CoV-2 that are more transmissible and virulent emerged raising doubts about their efficiency. Therefore, this work aims to explain possible molecular mechanisms responsible for the increased transmissibility and the increased rate of hospitalizations related to the new variants. A combination of theoretical methods was employed. Constant-pH Monte Carlo simulations were carried out to quantify the stability of several spike trimeric structures at different conformational states and the free energy of interactions between the receptor binding domain (RBD) and Angiotensin Converting Enzyme 2 (ACE2) for the most worrying variants. Electrostatic epitopes were mapped using the PROCEEDpKa method. These analyses showed that the increased virulence is more likely to be due to the improved stability to the S trimer in the opened state (the one in which the virus can interact with the cellular receptor ACE2) than due to alterations in the complexation RBD-ACE2, once the increased observed in the free energy values is small. Conversely, the South African variant (B.1.351), when compared with the wild type SARS-CoV-2, is much more stable in the opened state (either with one or two RBDs in the up position) than in the closed state (with the three RBDs in the down position). Such results contribute to the understanding of the natural history of disease and also to indicate possible strategies to both develop new therapeutic molecules and to adjust the vaccine doses for a higher production of B cells antibodies.

**Graphical Abstract:** 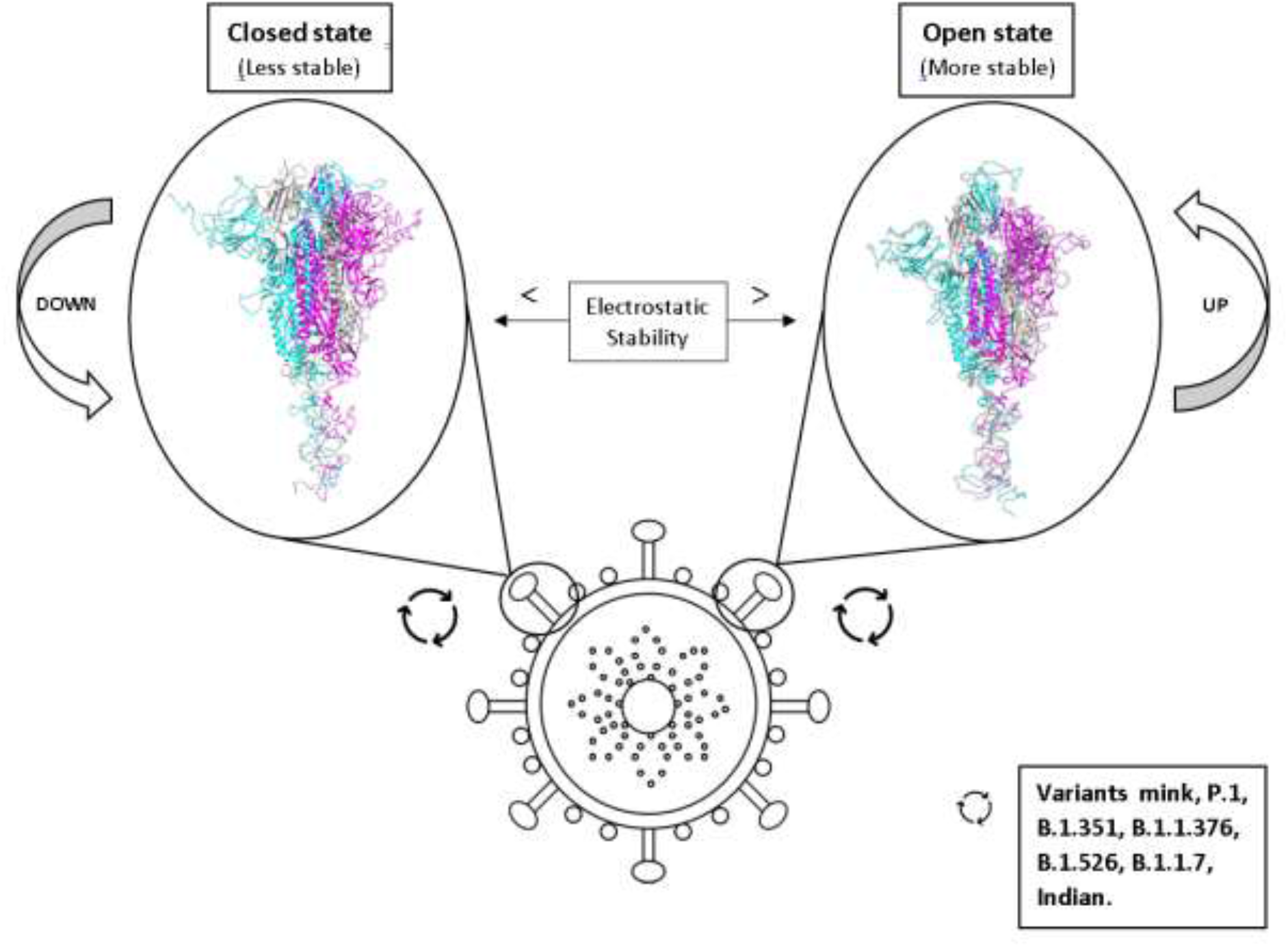

## 1. Introduction

SARS-CoV-2, the new beta coronavirus responsible for the COVID-19 pandemic, was able to infect more than one hundred million people and to kill more than two millions, approximately 110 times more than the 2009 H1N1 pandemic officially killed in one year (1,2). Even though the numbers are already quite alarming, the data is not being collected evenly, due to different coping strategies adopted worldwide. Moreover, the test results are not blindly reliable. The RT-PCR test, for example, considered to be trustworthy, was found to have high false-negative rates due to insufficient cellular material and different viral load kinetics of SARS-CoV-2 depending on the patient and sampling timing (3–5). The numbers of the COVID-19 deaths, as a result, are underestimated and far from reality. The 2009 H1N1 pandemic mortality, for example, was 15 times higher than the number shown by laboratory-confirmed results (2).

Both SARS-CoV-2 and SARS-CoV-1, the other coronavirus responsible for the 2003 Severe Acute Respiratory Syndrome pandemic in China, use a very similar mechanism to promote the fusion of viral and cellular membranes and to infect the human cell (6). They both interact with the ACE2 (Angiotensin Converting Enzyme 2) receptor through the Spike (S) glycoprotein receptor binding domain (which share 74% of sequence identity – see ref. (7) with a nearly identical binding conformation and similar affinities to promote the cell entry (6,8). Virus entry is also related to the presence of N-linked glycans around the post-fusion spike protein structure, which probably act in a protective role against host immune responses (9). Although the virus already has some immune escape mechanisms naturally, mutations can increase this escape, meaning that people who have already been infected could remain susceptible for reinfection, such as what happened in Manaus, Brazil (10). All viruses mutate as they replicate numerous times. In fact, although Coronaviruses have a relatively efficient proofreading mechanism (11), a large number of mutations have already been reported. For instance, the “Global Initiative on Sharing All Influenza Data” (GISAID) (12) repository has more than 804k (17 March 2021) genome assemblies available in their “official hCoV-19 Reference Sequence”. They adopted the high-quality genome sequence “hCoV-19/Wuhan/WIV04/2019”, isolated from a clinical sample at the Wuhan Jinyintan Hospital in Hubei Province on 30th December 2019, as the official reference sequence.

Typically, point mutations with a defective genome do not represent any health issue. However, mutations driven by an adaptive evolution are an advantage for the virus, and as such they represent a possible increased risk to human health raising doubts about the developed vaccines efficiency and antibodies therapies. The major concern arises from the knowledge that small differences in genetic material can substantially alter the properties of the viral proteins and offer extra advantages to viruses, such as higher ability to transmit or the capacity to escape the control of antibodies (whether produced due to a previous infection, received via intravenous administration or stimulated by vaccination) (13). Due to the rising fear of possible consequences that new variants may bring to the outcome of the pandemic, local outbreaks have been studied and some of the variants responsible for them are being classified as VOCs (variants of concern) by some health organizations from around the world, such as CDC (Centers for Disease Control and Prevention) and ECDC (European Centre for Disease Prevention and Control) (14,15).

To data, there are currently three principal variants considered VOC: a) 501Y.V1, also called VOC 202012/01 and B.1.1.7, b) B.1.351, also called 501Y.V2, and c) P.1 (15). All these three variants are of concern because of specific mutations that increased the virus’ transmissibility and, therefore, its harmful effects on the society. 501Y.V1, a variant detected in the United Kingdom approximately 50% more transmissible, carries eight mutations on the Spike glycoprotein homotrimer, but three of them are particularly worrisome – N501Y, meaning that residue number 501 had an asparagine replaced by a tyrosine, P681H and the deletion of residues 69 and 70 (16–18).

B.1.351, on the other hand, was first detected in South Africa, and has, apart from N501Y (included in the three VOC), the mutations K417T and E484K at the receptor binding domain (RBD), the same key mutations present in the P.1 lineage, detected in Brazil (19,15). Even though the impact of these mutations on the course of the pandemic is still partially unknown, different authors have elucidated some key aspects of the mutations mentioned. N501Y, for example, is located in the RBD and may increase ACE2 binding and transmissibility, while the infectivity rises by the deletion 69/70 (20,21). The other two mutations found in both B.1.351 and P.1, E484K and K417N, have a crucial role in viral escape preventing the neutralization by some antibodies (19,10). Although individually these mutations might generate changes in the trimeric’s properties, it is known that the combination of E484K, K417N, and N501Y mutations cause a greater conformational change in the RBD than N501Y or E484K alone (22). Another worrisome modification in B.1.351 and P.1 is the mutation D614G, once 614G variant, as well as ORF1ab 4715L, were proven to be related to higher fatality rates (23).

The main clinical aspects of COVID-19, unlike efficient treatment strategies, are well documented and known in the literature at least for the wildtype virus. The illness usually starts with nonspecific symptoms, such as fever, persistent cough, and fatigue (24). However, loss of smell and sense, delirium, skipped meals, and gastrointestinal symptoms, like abdominal pain and diarrhea, can also be considered to identify individuals infected with SARS-CoV-2 (25). In a more advanced stage, constituting the severe form of the disease, shortness of breath can also be observed, which usually leads to hospitalizations. However, the clinical aspects of COVID-19 caused by the new variants are still being elucidated.

Experimental data suggest that P.1 and B.1.351 are partially or totally resistant to antibodies developed for the treatment of COVID-19 and are inefficiently inhibited by the serum from convalescent individuals (26). Moreover, based on analysis from the “New and Emerging Respiratory Virus Threats Advisory Group” (27), from the United Kingdom, the infection caused by the B.1.1.7 variant is associated with a higher rate of hospitalization and death when compared to the wild type of SARS-CoV-2 (27). All these results suggest that the increased transmissibility and a potential antigenic escape are the reason why there is a resurgence of cases and why individuals previously infected with the wild type are only partially protected against the VOC (26,28).

Indeed, among several possible interdependent mechanisms that can increase viral transmissivity (e.g. increased viral shedding, longer interval of contagiousness, people mobility, wearing masks), a couple of them are directly related to mutations: a) an increased binding affinity between the viral adhesins with specific cell receptors (in this case, the RBD-ACE2 complexation, that defines the virus virulence and a possible increased infectivity), b) an increased environmental stability of the viral proteins and the virion particle, c) a higher availability of human cell receptors by either direct genetic conditions (e.g. the concentration of receptors ACE2 tends to be more elevated in men [29]) or due to commobidates (e.g. a lower stomachal pH in patients with Barrett’s esophagus induces a higher expression of ACE2 (30), d) an increased interaction with co-receptors, and e) immune evasion (31–34). Each of these individual factors and also their interplay have been discussed in the literature. For instance, there are works investigating other possible interactions for the spike protein to enter the cell (35,36), genetic factors affecting the viral virulence [e.g. the ACE2 polymorphisms can also influence the virus entry in the host cell and individual susceptibility (37), and the immune evasion (38–40).

The interaction between viral proteins and cell receptors can vary from virus to virus and also for different mutations (32,41). For the spike RBD, it has already been reported comparisons for its binding affinity with ACE2 for both SARS-CoV-1 and SARS-CoV-2 (42–45). However, it is still lacking data for the new variants of SARS-CoV-2 quantifying their impact on the virulence. In a landmark experimental work, Starr and co-authors partially addressed this question and performed a deep mutational scanning of SARS-CoV-2 RBD. They provided a handful set of information about the effect of *single* mutations on the binding, stability and expression (46). Yet, nothing is known about double or triple substitutions. Moreover, the spike trimer has to undergo a complex mechanism to make the RBD available for a proper binding with ACE2. Cryo-electron microscopy (CryoEM) structures of the SARS-CoV-1 spike trimer revealed that at least one chain of the homotrimer has to be at the “up” position (“open” conformation) as a prerequisite conformational state for the RBD-ACE2 interaction (47). At least a two-step “expose–dock-like” mechanism is needed to first allow the whole homotrimeric structure to perform all the conformational adjustments (first step of this complex process) before steric clashes (seen for the Spike “close” conformation) are removed to allow the complexation RBD-ACE2 to happen (second step) (43,47). Up to three receptors ACE2 can be bound one by one (48). Since several mutations present in the new variants occurred at the spike protein outside the RBD region (e.g. for the 501Y.V2 variant, L18F, D80A, D215G, R246I, D614G and A701V), it is expected that they have an impact on the “up” and “down” mechanism of the expose step. Also, the number of titratable amino acids involved in these mutations naturally suggests an electrostatic dependence. Nevertheless, the pH effects are somehow contradictory or, at least, not fully understanded. By one side, pH does not seem particularly important to trigger the conformational changes from the “down” to the “up” state (49). For example, no significant conformational changes were observed for CryoEM obtained at lower pH (pH 5.6) in comparison with neutral-pH (pH 7.2) for SARS-CoV-1 (49). By another side, Zhou et al. (2020) concluded that an immune evasion could be facilitated for SARS-CoV-2 by the low pH down conformation because of a pH-dependent refolded region located at the spike-interdomain interface – consisting of residues 824-858 – that exhibited structural modifications and RBD-mediated positioning of the trimer apex (50).

Following a previous work on the interactions of the RBDs of SARS-CoV-1 and SARS-CoV-2 spike proteins and the human cell receptor ACE2 (43), we investigated here the interactions of the RBD of the new most worrying variants (at the present) and other mutations with ACE2. By means of constant-pH biophysical simulation methods, the binding affinities between the RBDs of these variants with ACE2 were quantified at different pH regimes and the electrostatic epitopes of each case mapped and compared. For this analysis, the spike RBD was assumed to be totally ready for the interaction at the proper conformational state (i.e. only the second step of the “expose–dock-like” mechanism was studied). Another important aspect explored in this work was the electrostatic stability of the different possible conformational states of the trimer [all S chains at the “down” state (DDD), one chain at the “up” state and two others “down” (UDD) and two chains at the “up” state and one “down” (DUU)] for these new variants at all pH regimes. Such combined data can provide an answer for the increased transmissivity of the new variants observed in the clinical practice explaining the faster spread of them, the increased number of hospitalizations and a tendency to affect younger patients.

## 2. Materials and Methods

The field of virology is being widely explored and enriched by computational approaches, involving from bioinformatics tools and machine learning methods to biophysical simulations to understand the various aspects of viral functioning, including its immunology, pathogenesis, structural and molecular biology of the virus proteins (51–53). Complementing experimental studies, computational tools allow, for example, the understanding of molecular mechanisms of the virus, such as the comprehension of capsid proteins assembly (assembly intermediates are still difficult to be obtained by lab experiments), the quantification of the biomolecular interactions of the host-pathogen system, the prediction of conventional and electrostatic epitopes (EE), the understanding of increased virulence for different strains, and the development of specific molecular binders for diagnosis, treatment and prevention (43, 54–60).

Biophysical simulations of virus systems, based on computational molecular simulation methods such as Monte Carlo (MC) (61,62) and classical Molecular Dynamics (MD) (61,63) can take advantage of their long recorded success for probing the thermodynamic, dynamic, and interactive properties of biomolecules in pharmaceuticals (see Refs. (64,65), for reviews). Here, a fast constant-pH MC scheme (66,67) is applied to identify important residues for hostpathogen interactions and to clarify the intermolecular interactions involving the RBD of S proteins of SARS-CoV-1, 2 and the South African variant (B.1.351) and the electrostatic stabilities (68) of the spike homotrimers for the DDD, DUU and UDD conformational states at all solution pHs. Additionally, other recent variants like the Brazilian P.1 (alias of B.1.1.28.1), the Californian (B.1.1.376), the New York (B.1.526), the Indian double mutations E484Q and L452R (not classified yet) and the SARS-CoV-2 mink-associated variant strain (Y453F) were included in some of our analyses.

### 2.1. Molecular systems and their structural modeling

Several molecular systems were investigated in the present study employing the SARS-CoV-1, 2 and the new variants of the S RBD proteins with ACE2 (simulation set 1). Also, the electrostatic stability of whole spike homotrimeric protein at different conformational states were also studied (simulation set 2). A scheme of these two simulation sets is given in Figure 1. The three dimensional coordinates of these macromolecules necessary for all these simulations were obtained from different sources:

i. (a) the **SARS-CoV-1 S RBD wildtype (wt) protein (RBD1_wt_):** it was extracted from the RCSB Protein Data Bank (PDB) (69) where it was deposited with the PDB id 2AJF (chain E, resolution 2.9 Å, pH 7.5) and found complexed with ACE2 (chain A) – see Fig. 2.
ii. (b) the **SARS-CoV-2 S RBD wt protein (RBD2_wt_)**: for the sake of comparison with a previous work (43), we used the coordinates obtained from the comparative modeling of the protein three-dimensional structure built up at the SWISS-MODEL workspace (YP_009724390.1) based on the NCBI reference sequence NC_045512 (70). See ref. (43) for the details. The rootmean-square deviation (RMSD) of atomic positions between this modeled structure for the RBD of SARS-CoV-2 S wt protein and the available one for SARS-CoV-1 (PDB id 2AJF) is 0.64Å. A comparison with experimental SARS-CoV-2 S RBD structures deposited *after* this theoretical model was proposed (e.g. PDB ids 6W41 [resolution 3.08 Å, pH 4.6] and 6YM0 [resolution 4.36 Å, pH 8]) revealed similar RMSDs (0.51-0.56Å) to this one (0.64Å). This is an important feature showing that the sequence-structure relationship is robust enough to handle even mutations while still preserving the overall fold of the molecules. Previous works have also indicated this feature (71). Additional runs were performed with RBDs extracted from the PBD ids 6VSB (prefusion open state with one RBD at the up position) and 6VXX (close state). These are CryoEM coordinates obtained with resolution of 3.46 and 2.8Å, respectively. We shall refer to them as RBD2wt’_(6VSB)_ and RBD2wt’_(6VXX)_.
iii. (c) the **SARS-CoV-2 S RBD variants (RBD2_variant_):** the coordinates for all studied variants were modelled by the simple replacement of amino acids from the SARS-CoV-2 S RBD wt structure followed by an energy minimization using “UCSF Chimera 1.14” (72). Replaced amino acids are a) N501Y, K417N and E484K (19,46,73,74), for the South African (SA) variant (RBD2_SA_), b) K417T, E484K, N501Y (75–78), for the Brazilian (BR) P.1 (RBD2_BR_), c) Y453F (79) for the “mink” (RBD2_m_) strain, d) N501Y (40,75), for the UK B.1.1.7 variant (RBD2_UK_), e) E484K and N501Y (80), for the New York (NY) B.1.526 variant (RBD2_NY_), and f) L452R (81), for the Californian (CA) B.1.1.376 (RBD2_Ca_) strain. Note that both the SA B.1.351, the Brazilian (P.1) and the NY B.1.526 variants share two common mutations (E484K and N501Y). The third mutation at the RBD for the BR and SA variants is absent in the NY variant and occurs at K417 that is replaced by N and T, respectively for the BR and SA variants, and does not alter the electrostatic properties of the RBD. Both ASN and THR have the same physical chemical characteristic being polar with uncharged side chains indicating that these mutations are a natural adaptation that helps the virus. As a matter of fact, the E484 was also quite recently found in a double mutation (E484Q and L452R) in India (RBD2_I_).
iv. (d) the **SARS-CoV-2 S homotrimer wt protein (Strimer2_wt_)**: Structural data is now abundant for the spike proteins (or its parts) providing us with a rich level of information never seen before. To date, 151 experimental structures are available at the PDB with a diversity of resolution, conformational states and bound forms (apo and holo with different partners). They were solved by either X-ray crystallography or CryoEM. A good compilation of these available structures can be found at “Universal Protein Resource” (UniProt) under the id P0DTC2 (82). Several computer simulations were also performed expanding even more the information that can be extracted from these experimental structures (83–85). Despite this amazing work in such a short time, new variants are surging also in a highly competitive time frame, and key physical chemistry parameters as the solution pH were not explored in full details yet. From this ample source of available structural coordinates, we decided to use as input structures for our present calculations configurations extracted from the MD trajectories generated at Poma’s lab (84). This research group has characterized the structural and energetic differences between the DDD, DUU and UDD conformations of the SARS-CoV-2 spike wt trimer at pH 7. The advantages of this choice are: *i)* standardization of all physical chemical conditions used to obtain the structures at the three conformational states (different experimental structures were solved at distinct conditions) in the absence of any additional chemical (i.e. in a genuine electrolyte solution), ii) *all* structures from these MD trajectories were completed (no missing amino acids) and already minimized while the experimental structures have missing residues (in fact, the CryoEM structures contain several missing amino acids particularly at the RBD), *iii)* this set of configurations includes thermal structural fluctuations, *iv)* configurations extracted from different simulation replicas allow us to estimate the standard deviations in our own measurements. All these issues avoid the introduction of additional artifacts in our calculations. We also performed simulations with the experimental structure given by the PDB id 7A94 (CryoEM SARS-CoV-2 Spike hometrimer wt with one ACE2 bound, resolution 3.9Å, pH 8). Being this configuration (UDD) the most populated conformational state experimentally observed among the bound ones (48), this allows us to test an experimental structure and how the binding of ACE2 changes the stability of the complex S-ACE2.
v. the **SARS-CoV-2 S homotrimer SA variant (Strimer2_SA_)**: assuming valid the robust sequence-structure relationship to preserve the overall fold of the molecules upon point mutations (as mentioned above), the coordinates for all studied variants were modelled as described above for item (c) by the simple replacement of amino acids from the SARS-CoV-2 S wt trimer structures at the corresponding three different conformational states. Replaced amino acids are for the SA B.1.351 variant, D614G, N501Y, K417N, E484K, L18F, D80A, D215G, R246I, A701V (19,46,73,74).
vi. the **SARS-CoV-1 S homotrimer wt protein (Strimer1_wt_)**: as done for the SARS-CoV-2 B.1.351 variants, deletions and amino acids substitutions were performed in the SARS-CoV-2 wt homotrimer following the sequence of the SARS-CoV-1 S protein as given by Uniprot id P59594 (82). This introduces an additional approximation in our outcomes whose effect is assumed to be small due to the high identity (at the sequence level) between them and the strong sequence-structure relationship mentioned above. Additional calculations were performed with the experimental structures given by PDB ids 6ACC (DDD conformational state) and 6ACD (UDD conformational state).

**Figure 1.**
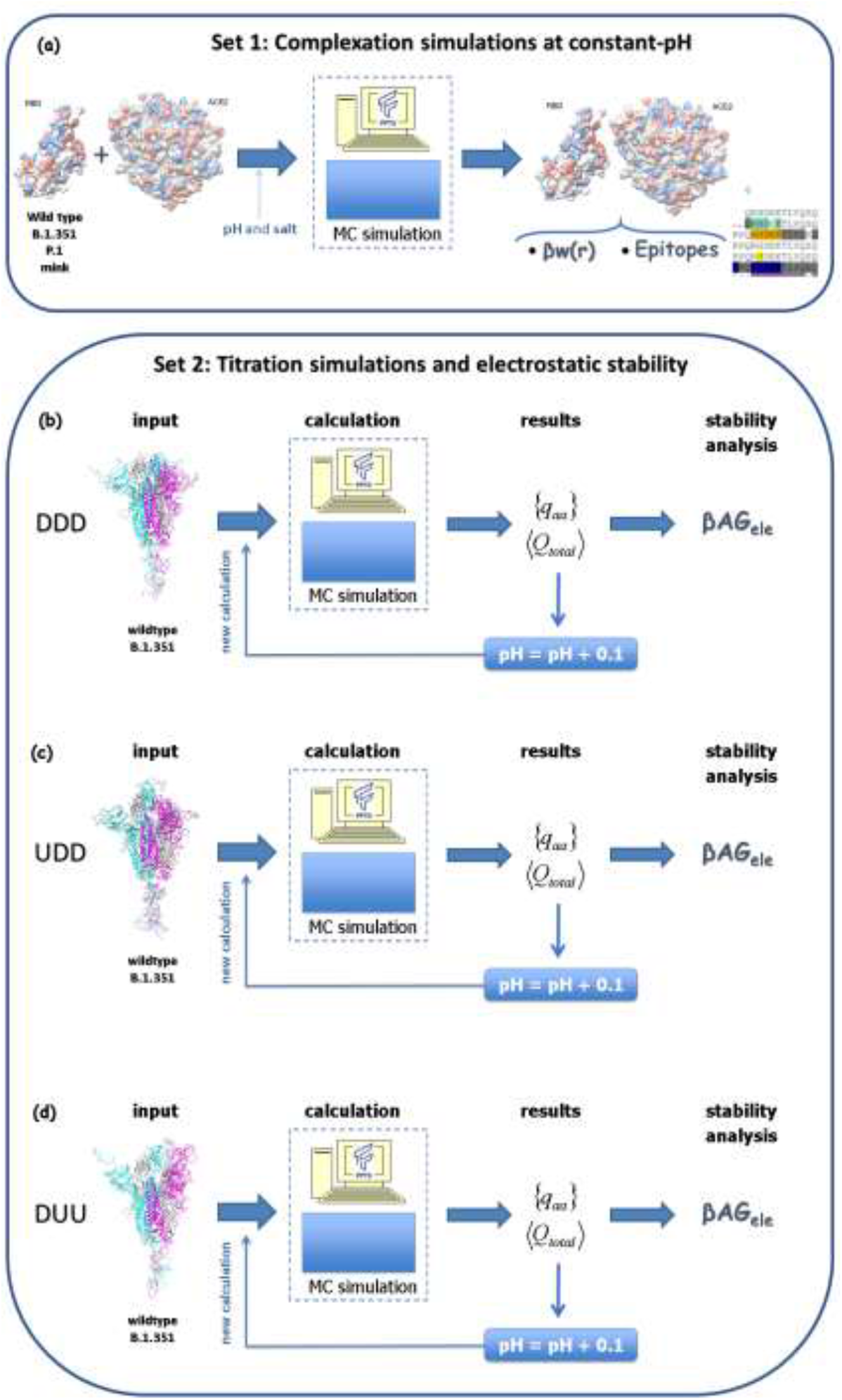
Schematic representation for the two simulation sets of this work. Simulation set 1 (a) represents complexation simulations at constant-pH where the interaction between the S RBD protein of SARS-CoV-1, 2 and the new variants (B.1.351, P.1 and the mink variant) with receptor ACE2 were investigated. Simulation set 2, consisting of (b), (c) and (d), represents simulations used to estimate the electrostatic stability of whole spike homotrimeric proteins at different conformational states (DDD, UDD and DUU, respectively) for all solution pHs. Spike wt proteins from SARS-CoV-1, 2 and its B.1.351 were investigated.

**Figure 2.**
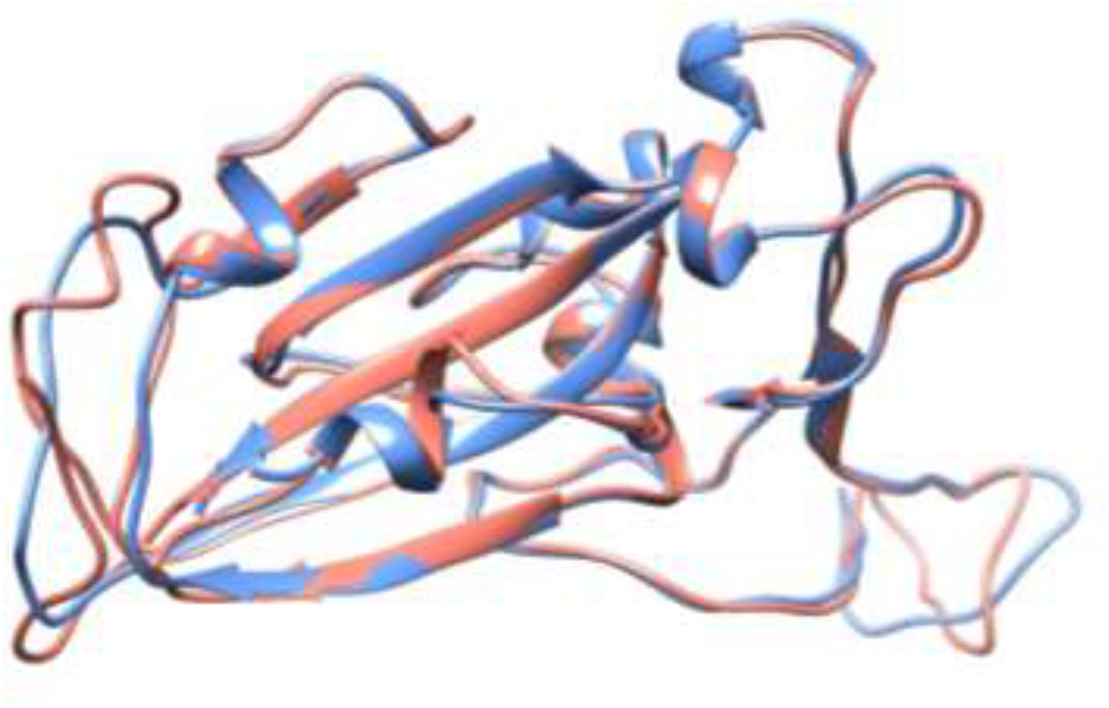
Crystal structure of the SARS-CoV-1 S RBD (PDB id 2AJF, chain E) and the modeled SARS-CoV-2 S RBD (wildtype). See text for details regarding the modeling aspects. These macromolecules are shown, respectively, in blue and red in a ribbon representation. The RMSD between these structures is equal to 0.64Å.

**Figure 2.**
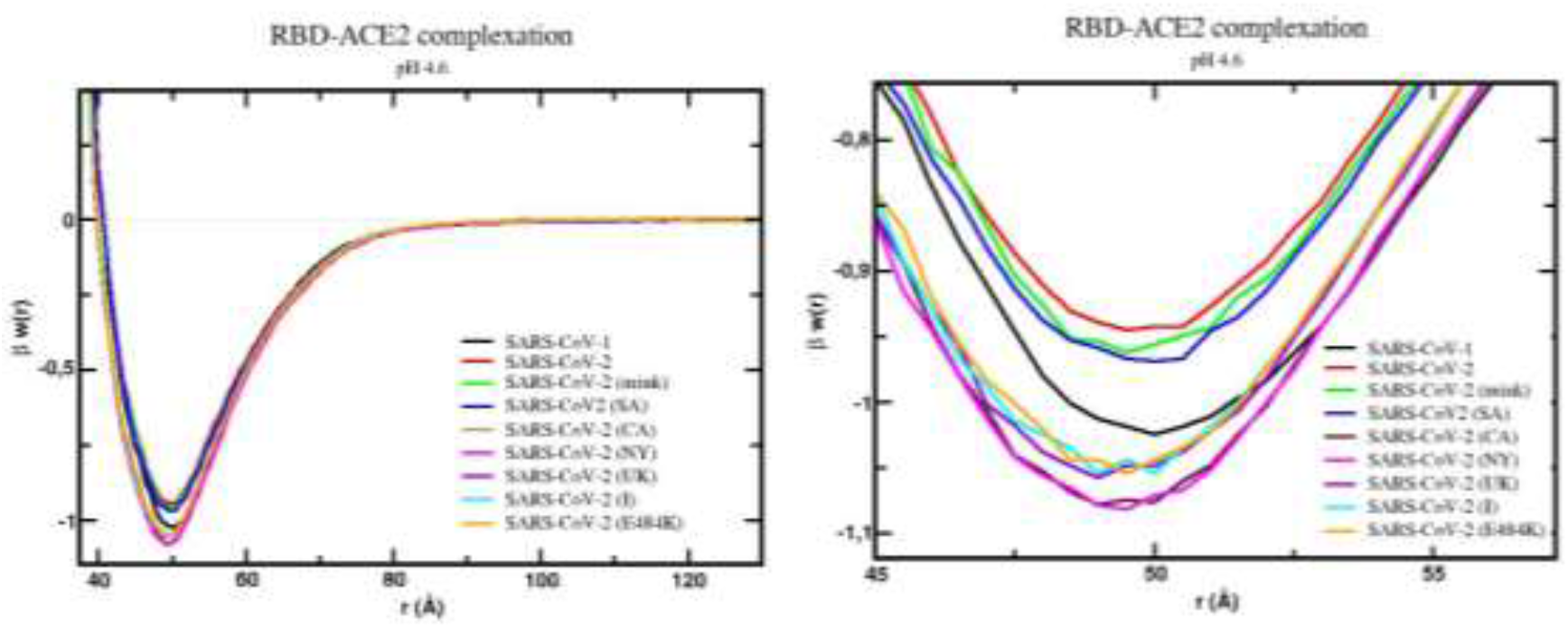
Free energy profiles for the interaction of RBD proteins with the cellular receptor ACE2. The simulated free energy of interactions [**β**ι*w*(*r*)| between the centers of the RBD proteins from SARS-CoV-1, SARS-CoV-2 (wt), SARS-CoV-2 (mink), SARS-CoV-2 (SA), SARS-CoV-2 (CA), SARS-CoV-2 (NY), SARS-CoV-2 (UK), SARS-CoV-2 (I), and SARS-CoV-2 (E484K) and the cellular receptor ACE2 are given at pH 4.6. The source of the three dimensional structures of these proteins are explained in the text and referred as RBD1_wt_, RBD2_wt,_, RBD2_m_, RBD2_SA_, RBD2_CA_, RBD2_NY_, RBD2_UK_, RBD2_I_, and RBD2_E484K_, respectively. Salt concentration was fixed at 150 mM. Data for the complexes with the wildtype proteins (RBD1_wt_-ACE2 and RBD2_wt_-ACE2) was given before (Giron *et al.*, 2020). Simulations started with the two molecules placed at random orientation and separation distance. Results for SARS-CoV-1, SARS-CoV-2 (wt), SARS-CoV-2 (mink), SARS-CoV-2 (SA), SARS-CoV-2 (CA), SARS-CoV-2 (NY), SARS-CoV-2 (UK), SARS-CoV-2 (I), and SARS-CoV-2 (E484K) are shown as black, red, green, blue, dark purple, pink, light purple, cyan, and orange continuous lines, respectively. (a) *Left panel:* Full plot. (b) *Right panel:* The well depth region of the β*w*(*r*) for each studied complex.

As indicated above between parenthesis, we shall refer to these systems (a)-(f) as RBD1_wt_, RBD2_wt_ (and RBD2_wt’_), RBD2_variant_ (variant = SA, BR, CA, NY, UK, I or m), Strimer2_wt_, Strimer2_SA_, and Strimer1_wt_, respectively. Calculations performed with a different input structure will have their PDB ids included as a subscript (e.g. Strimer1_wt(6ACC)_ for the SARS-CoV-1 spike wt protein using the PDB id 6ACC). The receptor ACE2 needed for all simulations of set 1 was obtained from the PDB id 2AJF (chain A) as in a previous work (43).

All PDB files were edited before the calculations. Missing regions in these proteins were built up using the “UCSF Chimera 1.14” interface (72) of the program “Modeller” with default parameters (86). Water molecules and hetero atoms were completely removed from all used files. Glycosylation sites were not included (they are also often truncated in the experiments) due to the incompatibility of highly flexible molecules with a rigid protein model and all the uncertainties and arbitrariness involved in it (87). Cysteines involved in sulphur bridges were fixed with GLYCAM web tools (88). The “UCSF Chimera 1.14” package (72) was employed for all molecular visualizations and representations too. Some images were generated by CoV3D (89). When appropriate, it is indicated at the figures captions. For some analysis, it was necessary to determine structural interfaces. This was done with the online server “PDBePISa” (90) with default options.

The linear sequences of the SARS-CoV-1 and 2 S proteins are available in UniProt with the ids P59594 and P0DTC2 for SARS-CoV-1, 2 (wt), and B.1.351, respectively. The S1 subunit that binds the virion to the cell membrane receptor is the cleaved chain between residues 14 and 667 for SARS-CoV-1, 13 and 685 for SARS-CoV-2. To our knowledge, there is no information yet if the mutations have affected the cleaved region for SARS-CoV-2. Alignments of pairwise sequences were obtained by the EMBOSS Needle server (91) with default settings. They are shown in Figures S1-3. The identity and similarity between SARS-CoV-1 and 2 wt are 64.2% and 78.6%, respectively. The RBD corresponds to positions 306-527 and 319-541 for SARS-CoV-1 and 2, respectively. Both identity (I) and similarity (S) are higher for this specific structural region (I = 73.1 % and S = 82.1%). On the other hand, the identity and similarity between SARS-CoV-1 and the B.1.351 variant are 68.2% and 75.5%, respectively. The identity and similarity between SARS-CoV-2 wt and the B.1.351 variant are 98.4% and 99.0%, respectively. This comparison shows that the B.1.351 variant of SARS-CoV-2 has an identity for the RBD slightly closer to SARS-CoV-1 than SARS-CoV-2 wt (68.2% against 64.2%).

### 2.2. Molecular simulations

Despite the wide availability of molecular simulation models at different scales (64, 92–95), the so-called coarse-grained (CG) models are the most cost-effective ones, due to *i)* the large number of atoms involved in each of the studied systems (e.g. the SARS-CoV-2 S wt homotrimer has 3363 amino acids), *ii)* the electrostatic coupling between a large number of titratable groups in these macromolecular structures (e.g. 246 groups per chain of the SARS-CoV-2 wt homotrimer), *iii)* the need to repeat the calculations both at several different physical chemical conditions (140 different pH conditions per system), for different protein conformations (e.g. 10 input structures for each conformational state of the homotrimer) and several viral protein systems (more than 7 systems as described below, see item 2.1), *iv)* the estimation of the free energy of interactions based on a histogram method where the statistics on each histogram bin requires longer simulation runs for a proper sampling, and *v)* the number of simulation replicates to guarantee their numerical convergence. These simplified computer models allow a lower computational cost for the exploration of the main physical characteristics of a system with a small number of parameters (54,97,98). Though it may not be so explicit, all these calculations required extensive computational resources.

A simple CG for protein-protein interactions has been devised using a rigid body description of the macromolecules, and successfully applied to study several different biomolecular systems (54,97–100). The main features of this model is the inclusion of a fast and accurate description of the pH effects by means of the fast proton titration scheme – FPTS (66,67,95,96). Ideally, the coupling between the proton equilibria and conformational changes should be described by a constant-pH molecular dynamics (CpH MD) scheme. Nevertheless, convergence specially of the electrostatic properties is still a critical issue of such methods for a single and small protein (101–104). The cpu costs are prohibitive to study all the systems and conditions mentioned above even for fast CpH MD approaches as our OPEP6 (104). Conversely, an interesting observation is that key dynamical properties have a significant relation with electrostatic property variations as recently demonstrated for flaviviruses (41). Therefore, we followed the constant-pH MC (CpH MC) strategy as used before for hostpathogen interactions (43,59). Other models are also available in the literature. For instance, Yu et al. (2021) are currently developing a multiscale CG model of the SARS-CoV-2 virion, which helps with the molecular simulations an amazing description of the SARS-CoV-2 virions because of the model’s ability to adapt as new information about the molecule is released. However, pH effects are not fully included in the model as it is done in our present work.

For the present CpH CG model, each group of atoms defining an amino acid is converted into a single charged Lennard-Jones (LJ) sphere of valence zi (a function of the pH of the solution) and radius (R_i_) – the values for each class of amino acid being taken from the work of Persson *et al.* (2010). The centers of mass of the spheres created are used to arrange them accordingly with their experimental three-dimensional structures (as specified above). In order to obtain the valences and allow the variation of the amino acids depending on the pH during the simulation, FPTS was employed, whose physicochemical basis and explanation can be found in previous publications (66,67,96,101).

For all simulations of set 1 (i.e. for the complexation RBD-ACE2 – see Fig 1), two proteins (here, the RBD and the ACE2) are placed in an open cylinder simulation box to allow for forward and backward translations in one axis combined with rotational movements in any direction. As for the simulation data, the static dielectric constant of the medium (ε_s_) was 78.7 to mimic an aqueous solution (assuming a temperature of 298 K), the radius (r_cyl_) of the simulation cell used was 150Å and height (l_cyl_) was 200Å. Furthermore, the salt particles and the added counter-ions were represented using an electrostatic screening term as follows: for two ionizable amino acids *i* and *j,* the screening is given by [exp(-κr_ij_)], where κ is the modified inverse Debye length, and r_ij_ is the distance between particles (54,97,98,101). For simulations of set 2 (i.e. the stability of different homotrimers), only one macromolecule was included in the simulation cell.

The electrostatic interactions [u_el_(r_ij_)] between any two ionizable amino acids of valences *z_i_* and *z_j_* are given by:

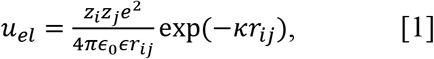

where *e* (the elementary charge) is 1.602×10^-10^C and *ε_0_* is the dielectric constant of the vacuum (*ε_0_* = 8.854×10^-12^C^2^/Nm^2^). Except for the ionizable amino acids charges – which were defined by the FPTS (66,67) – all the others were fixed neutral and kept constant during all simulation runs. Further details can be found on refs. (54,97,98,100).

Hydrophobic effect, van der Waals interactions and excluded volume repulsion can also affect protein-protein interactions (54,98,99,105). A simple way to at least incorporate the main contributions of these interactions is by means of a LJ term [u_vdw_(r_ij_)] between the amino acids (54,96,98). For any two amino acids (either charged or not), the calculation for the LJ term is given by

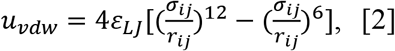

where *ε_LJ_* regulates the intensity of the attractive forces in the system (54,97,98), and σ_ij_ (= R_i_ + R_j_) is the distance between two amino acids, represented by *i* and *j*, when they are in contact. The choice of *ε_LJ_* is somehow arbitrary, although it has been used in many works an universal value of 0.124 kJ/mol (96–98,106), corresponding to a Hamaker constant of ca. 9k_B_T (where k_B_ = 1.380 × 10^-23^ m^2^ kgs^-2^K^-1^ is the Boltzmann constant, and *T* the temperature, in Kelvin) for amino acid pairs (98,99,107). The direct impact is that the calculated magnitude of the free energy of interactions can be either under or overestimated as we discussed before (43). For instance, the absolute numbers might be shifted when compared with results obtained using other force field descriptions. In principle, when experimental second virial coefficients are known for exactly the same physical chemical conditions and system, a proper calibration of *ε_LJ_* can be obtained for this very specific situation. This is not the case for these spike proteins. Alternatively, results can be interpreted in relative terms, comparing the measurements between similar systems and conditions. This solves the possible problem of ambiguity in the interpretation of the obtained data.

The size of the amino acids beads also affects the LJ contributions. For the sake of consistency with the adopted *ε_LJ_* value, all *R_i_*′s were taken from ref. Persson *et al.* (2010). Each amino acid has a specific value of *R_i_* (e.g. R_TYR_=4.1Å, R_GLU_=3.8Å. For this pair, σ_ij_= R_TYR_+R_GLU_=7.9Å) which allows the description of mostly macromolecular hydrophobic moments (108). Therefore, the simulations should correctly generate the docking orientation at short separation distances (54).

By bringing equations 1 and 2 together, one can obtain the total interaction energy of the system (whether charged or neutral) for a given configuration [U({r_k_})]:

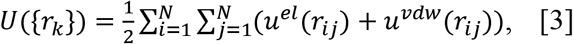

where {r_k_} is the position of the amino acids and N is the total number.

The results were obtained using the Metropolis MC sampling performed with physiological ionic strength (NaCl at 150 mM) at different pH regimes. For the complexation study described in the simulation set 1, following a previous study (43), the pHs 7.0 and 4.6 were chosen, respectively, due to the need to understand the behavior of the system in a physiologic pH levels condition and the acidic pH of the endosomal environment. This is a reasonable choice to keep the general features of the present study despite the fact that the precise value of the pH in human cells can be slightly different from these numbers (109). For all simulations from set 2, pH was varied from 0 to 14 with an increment of 0.1 units of pH to explore all possible pH conditions. This pH range was also used for the epitopes mapping by the PROCEEDpKa method (54) – see below. As indicated by Figure 1, the main outcomes of these CpH simulations were the averaged total protein charge (<Q_total_>), the averaged partial charge of each titratable group ({<q_aa_>}), the averaged protein dipole moment (<μ>). These quantities are also conveniently expressed in units of the elementary charge: the averaged total protein charge number (Z=<Q_total_>/e), the averaged valence of each titratable group (z_aa_=<q_aa_>/e), the averaged protein dipole number moment 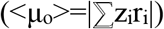. Free energy of interactions [**β**w(r)] were estimated from radial distribution functions [g(r)] between the centers of the proteins [**β**w(r)=-ln g(r), where **β** = 1/k_B_T]. *r* is the separation distance between the centers.

After the steps of preparing the molecular systems for the simulations described above, equilibration and production runs were performed. Even invoking all the approximations in the CG model, this demanded high computational resources due to: i) the large number of titratable groups involved in the system with strong electrostatic coupling; ii) the free energy barriers of the systems; iii) the need to fill all histogram bins used for the calculation of g(r) during sampling; iv) longer runs to the decrease statistical noises in the **β**w(r) data (54,97,98). All simulations from set 1, required at least 3.0 10^9^ MC steps at the production phase. Simulations from set 2 could be well performed with 10^8^ MC steps at the production phase. Standard deviations were estimated by the use of at least three replicates per simulated system. Some systems were further explored with additional replicates.

### 2.3. Electrostatic epitopes determined by the PROCEEDpKa method

Electrostatic properties are well known to be of great importance in biomolecular interactions. Indeed, they strongly depend on the spatial distribution of intra and intermolecular charges, environmental conditions, such as pH or salt concentration that can vary significantly in different cellular compartments (54,67). Due to its intrinsic long range characteristics and the electrostatic coupling between ionizable residues, key amino acids responsible for the hostpathogen interaction can include groups outside the classical view of the epitope-paratope interface. Such broader definition including inner titratable residues that can also take part in the interplay of interactions has been called “electrostatic epitopes” (EE). They can be efficiently mapped by a computational strategy called “PROCEEDpKa” (PRediction Of electrostatic Epitopes basedED on pKa shifts) (54). This allows the identification of all ionizable residues of macromolecules that do drive the biomolecular interactions. The difference between the numbers of classical and electrostatic epitopes will be higher as stronger is the electrostatic coupling between superficial residues with the inner ones. Among other advantages (see Ref. 54), “PROCEEDpKa” includes the pH and ionic strength dependence that can dramatically affect the complexation process of the host-pathogen interactions. This is particularly important for the antibody-antigen interface that has a peculiar electrostatic pattern richer in titratable amino acids.

### 2.4. Electrostatic stability

The electrostatic stability of the different spike homotrimers (Strimer1_wt_, Strimer2_wt_, and Strimer2_SA_) and their conformational states (DDD, UDD and DUU) as a function of solution pH were estimated by means of the electrostatic free energy (ΔG_elec_). This physical quantity was calculated in terms of Coulombic contributions arising from the individual titratable groups for a given protein structure in a specific conformation and physical chemical conditions (68). As proposed by Ibarra-Molero and coauthors (110), ΔG_elec_ (in kJ/mol) can be given as

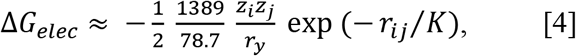

where *r_ij_* is the separation distance between the ionizable sites *i* and *j* as defined by the spike homotrimer conformation. All charges numbers *z_k_* are the averaged ones obtained from the titration studies, i.e. *z_k_*=<q_aa_>/e for a given solution pH, protein system and conformational state. κ was fixed at 7.86 Å to correspond to an electrolyte solution 1:1 with 150 mM NaCl.

## 3. Results

### 3.1. Free energy of interactions of SARS Spike RBD proteins and ACE2

An essential step in the process of cell invasion by a virus is the interaction with a cell receptor. The receptor used by SARS-CoV-1 has been known since 2003: the angiotensin converting enzyme II, or ACE2 (111). Due to the great similarity between the viral proteins, the same receptor has been found to be used by the virus responsible for the COVID-19 pandemic (112). This has been repeatedly confirmed by different studies (6,37,43,112,113). Modifications in this molecular region (which can be visualized in Fig. S.4), either by mutations in the Spike protein or by alterations in the receptor itself, can change the free energy of the interaction and result in higher or lower affinity. This is easily seen for some mutations. For example, the mutation E484K presented at least in three new variants (B1.1.351, P.1 and B.1.526) has an acid residue (GLU) replaced by a basic one (LYS). This substitution changes the physical chemical nature of the residue 484 and implies a complete inversion of its electrostatic interactions. GLU can have its electrical charge (in elementary units) varying from −1 (when fully deprotonated) to 0 (when fully protonated) while LYS has a zero charge number when deprotonated, and is positively charged (+1) when protonated. All neighbouring ionizable groups might also be affected by this inversion. This kind of evolutive viral signature suggests that the virus is using the electrostatic properties to increase its virulence. In fact, when comparing the main simulated physical chemical quantities of the RBD proteins, we can clearly see the effect of the substitutions of the amino acids both at the protein charge number level (varying from +2.1 to +4.1, see table S.1) and the dipole number moment level (varying from 31 to 90, see table S.1). These results given between parenthesis were obtained by the FPTS at pH 7.0 and are summarized in Table S.1 for all studied variants. Data for pH 4.6 is included in this table too. Being the ACE2 receptor negatively charged at pH 7, the attractive charge-charge interaction will be stronger for most of the variants which favours the RBD-ACE2 complexation for them. The differences in the dipole moment numbers reveal that the binding orientation can also be altered for some systems.

This preliminary and simplest physical chemical analysis was investigated with more quantitative details of the complexation process. As a continuation of a previous study on the molecular interactions of SARS-CoV-1 and 2 wt S RBDs done at the beginning of the pandemic (43), we investigated now the binding association of the S RBD proteins new variants to ACE2. The RBD was assumed to be exposed and ready for the dock phase (i.e. the RBDs were out of the homotrimeric S protein assuming that the missed parts of the whole trimer do not interfere with the binding as suggested by the available crystallographic data (43). The mink, SA (B.1.351), the BR (P.1), the UK (B.1.1.7), CA (B.1.427/B.1.429), the NY (B.1.526), the India RBDs complexations with ACE2 were studied by means of our CpH MC proteinprotein simulations at pH 4.6. The wild type with a single E484K mutation was also included in this study. Free energy of interactions [βw(r)] calculated by means of potential of mean forces sampled during the simulation runs for these systems are shown in Figure 2 at physiological salt concentration. The estimated maximum standard deviations obtained by the comparison of results from at least three replicates are 0.01 for βw(r).

Simulations confirmed the binding of all RBDs to ACE2 as can be seen by the negative values of βw(r) around 50Å. The SARS-CoV-1 S RBD protein has the strongest tendency to bind to ACE2, as can it was previously reported in some theoretical and experimental studies (43,114,115). Yet, this is not a consensus in the literature (6,116,117). Other works have reported an opposite behaviour where the SARS-CoV-2 S wt RBD has the strongest affinity (118). This discrepancy can have different sources (e.g. use of other structural coordinates, molecular dynamical runs not long enough to sample larger structural fluctuations, glycosylation, presence of other monomers of the ACE2, use of the homotrimer instead of a single RBD, constant-charge versus constant-pH simulations, different experimental assays, other physical chemical conditions, etc.) (116). In fact, calculations performed with RBDs coordinates extracted from CryoEM structures will suffer from the lack of coordinates for this critically important region. For instance, the prefusion S homotrimer with a single RBD up as given by the PDB id 6SVB (chain A) has several missing amino acids at the RBD (INT....KVGGN....LFRKSNLKPFERDISTEIYQAGSTPCNGVEGFNCYF....NG....).

Although they can be fully reconstructed by homology modeling, it becomes difficult to precisely identify if the final model provides more reliable coordinates for the complexation studies than the one builded-up using the SARS-CoV-1 RBD as a template (with less unknown coordinates). Testing RBDs extracted from the CryoEM structures (completed with homology modeling) can result in binding affinities equivalent to what was measured for SARS-CoV-1 or slightly superior – see Figure S.5. As expected and also reported before (97,101), there is a dependence on the used structural coordinates for the protein-protein calculations. A different set of coordinates for ACE2 could also have an effect together with the possible structural rearrangements during the binding. Constant-charge MD simulations would not be a definitive answer as indicated by a previous study that also predicted a higher affinity of SARS-CoV-1 RBD–ACE2 binding (115). An ideal solution would be the possibility of applying CpH MC simulations to all the variants which are still not feasible at the present. Instead, we opted to use in this work the RBD modelled as before (43) and focus on the aspects given by the effects of different substitutions of the amino acids which is the main aim of the present study. It is a consensus from different studies that electrostatic interactions drive the RBD-ACE2 complexation in both cases (43,116) which means that CpH models (as used in this work) even with other approximations should better capture the main physics of the system.

Moreover, despite the good precision in the protein-protein simulations [0.01 units of βw(r) as mentioned above], the differences between the potential depths for each of these proteins sequences is small. Taking into account all the intrinsic approximations assumed in this work (including possible fluctuations due to structural dynamics), we think it is safer to conclude that these outcomes are indicating tendencies given by the differences in the linear sequences. Therefore, this analysis showed a slight tendency toward a higher affinity for ACE2 by all studied new variants. The highest affinity was found for the RBD2_NY_. In a crescent order, we have RBD2_wt_ < RBD2_mink_ < RBD2_SA_ and RBD2_BR_ < RBD2_I_ < RBD2_UK_ < RBD2_CA_ < RBD2_NY_. The BR variant P.1 (data not included in Figure 2 because it was on the top of the plot of the SA case) behaves identically to the SA variant (B.1.351) due to the presence of the same key mutations in both of them – E484K and N501Y at the receptor binding motif. The mutations K417N/T do not make any difference for these two variants in terms of their binding affinities at least that can be captured by our CpH CG model. In fact, these substitutions on K417 might work to inhibit the ACE2-affinity (46). It can also be seen the effect of the absence of the mutation K417 in the NY variant. This enhances the RBD-ACE2 affinity specially when compared with the two other variants (SA and BR) that share the same substitutions but have either K417N or K417T to decrease it. The key mutations at the RBD region are illustrated in Figure S.6. The increased affinity between RBD2_UK_ and ACE2 has recently been seen in another computational approach (119). This result also revealed that the binding affinity of these new variants are approaching and enhancing the affinity measured for SARS-CoV-1 that was more virulent.

Table 1 summarized the main theoretical results together with experimental data for single mutations. For the sake of clarity, mutations are also listed in this table. Although the experimental available data was not obtained exactly under the same conditions, it can be seen that they are qualitatively similar to the theoretical data. The presence of more than one mutation does not seem additive. For this set of studied cases, both N501Y and Y453F are mutations that experimentally resulted in the highest RBD-ACE2 affinities (46). The theoretical predictions confirm that the N501Y mutation increases this affinity. The difference between this strain with the NY and CA variants seen in our calculations are within the estimated errors (0.01). Conversely, the Y453F mutation is equivalent to the wild type in the theoretical predictions when the estimated errors are taken into account. Despite quantitative discrepancies, it is clear that the new variants have a tendency to be more virulent than the wild type SARS-CoV-2 due to the molecular properties of their more evolutively adapted RBDs.

**Table 1:** Estimated binding affinities for RBD-ACE2 interactions. The theoretical predicted binding tendencies was calculated taken the differences between the minima for **β***w*(*r*) of the wild type with the simulated value for a given variant at pH 4.6. (*) Data from ref. 46. See text for other details.

| **RBD structure** | **mutation(s)** | **Predicted binding tendencies (pH 4.6)** | **Experimental relative binding affinity for each single mutation<sup>(\*)</sup>** |
| --- | --- | --- | --- |
| RBD2<sub>mink</sub> | Y453F | 0.01 | 0.25 |
| RBD2<sub>SA</sub> | K417N<br>E484K<br>N501Y | 0.02 | -0.45<br>0.06<br>0.24 |
| RBD2<sub>BR</sub> | K417T<br>E484K<br>N501Y | 0.02 | -0.26<br>0.06<br>0.24 |
| RBD2<sub>CA</sub> | L452R | 0.12 | 0.02 |
| RBD2<sub>NY</sub> | E484K<br>N501Y | 0.13 | 0.06<br>0.24 |
| RBD2<sub>UK</sub> | N501Y | 0.11 | 0.24 |
| RBD2<sub>I</sub> | L452R<br>E484Q | 0.10 | 0.02<br>0.03 |
| RBD2<sub>E484K</sub> | E484K | 0.10 | 0.06 |

### 3.2. Estimated antigenic regions by the PROCEEDp*Ka* method

In this timely research field that is evolving so fast with new variants frequently appearing with an increased rate, it becomes challenging to catch up such “storms” of mutations and run calculations for all possible new cases. For this reason, we selected the SA variant (B.1.351) to be investigated in more detail during the last phases of the present study. An important issue is now to understand what amino acids of its RBD are the relevant ones for the complexation mechanism. By one side, these epitopes can be used to design peptides for vaccines. By another, they contribute to the characterization of the binding modes indicating interesting potential targets for specific therapeutic binders.

In order to determine the EE of the S RBD proteins of SARS-CoV-1, SARS-CoV-2 wt, and the SA variant (B.1.351) of SARS-CoV-2 for the ACE2 complex, the PROCEEDpKa method (54) was used. This method uses pKa shifts to identify the key amino acids responsible for a host-pathogen association. It is rooted in the physical chemical fact that the presence of an electric charge (a fixed charge or the instant charge of another ionizable residue at a given protonation state) can perturb the acid-base equilibrium of a titratable group. Consequently, identifying the pKa shifts is a practical means to probe intermolecular interactions as before demonstrated (120). The pKas were measured both from the theoretical titrations for the isolated RBDs and during computer simulations of a protein-protein complexation by the CpH MC scheme (simulations set 1).

The main questions to be addressed here are to determine whether the mentioned proteins share a common binding region when interacting to ACE2, and whether there is a change in the number of amino acids involved on it (which can indicate a more or less specific association). The found EE during the simulations were mapped at the sequence level to allow a direct comparison. These results are shown in Figure 3. In this Figure, the primary sequences of the RBDs of SARS-CoV-1, SARS-CoV-2 wt and 2’ (representing the SA variant) are superimposed. Amino acids identified as EE as classified by the PROCEEDpKa method are shown in blue. Although the general patterns observed for these three viral proteins are relatively similar, some differences can be seen both in the number of perturbed amino acids and their location. These differences show how the electrostatic theoretical method is sensitive to the different sequences as seen previously (54). Indeed, it was noted before that SARS-CoV-1 and 2 (wt) are antigenically different (118). The number of ionizable residues involved in the intermolecular interactions between SARS-CoV-1 S RBD and ACE2, SARS-CoV-2 S wt RBD and ACE2, and SARS-CoV-2’ (RBD2_SA_) and ACE2 pairs increased from 30 to 40 and from 40 to 43, respectively, with a high number of common EE. Qualitatively, the theoretical predicted antigency patterns for SARS-CoV-1 and 2 (wt) are the same from another work published laterly reporting 17 and 21 epitopes, respectively (118). Other theoretical methods also support this behaviour (115). The quantitative differences are due to the fact that the EE includes amino acids more internalized in the structure that are electrostatically coupled with the superficial ones. As a matter of fact, they can have an important role in the complexation together with the superficial residues (54).

**Figure 3.**
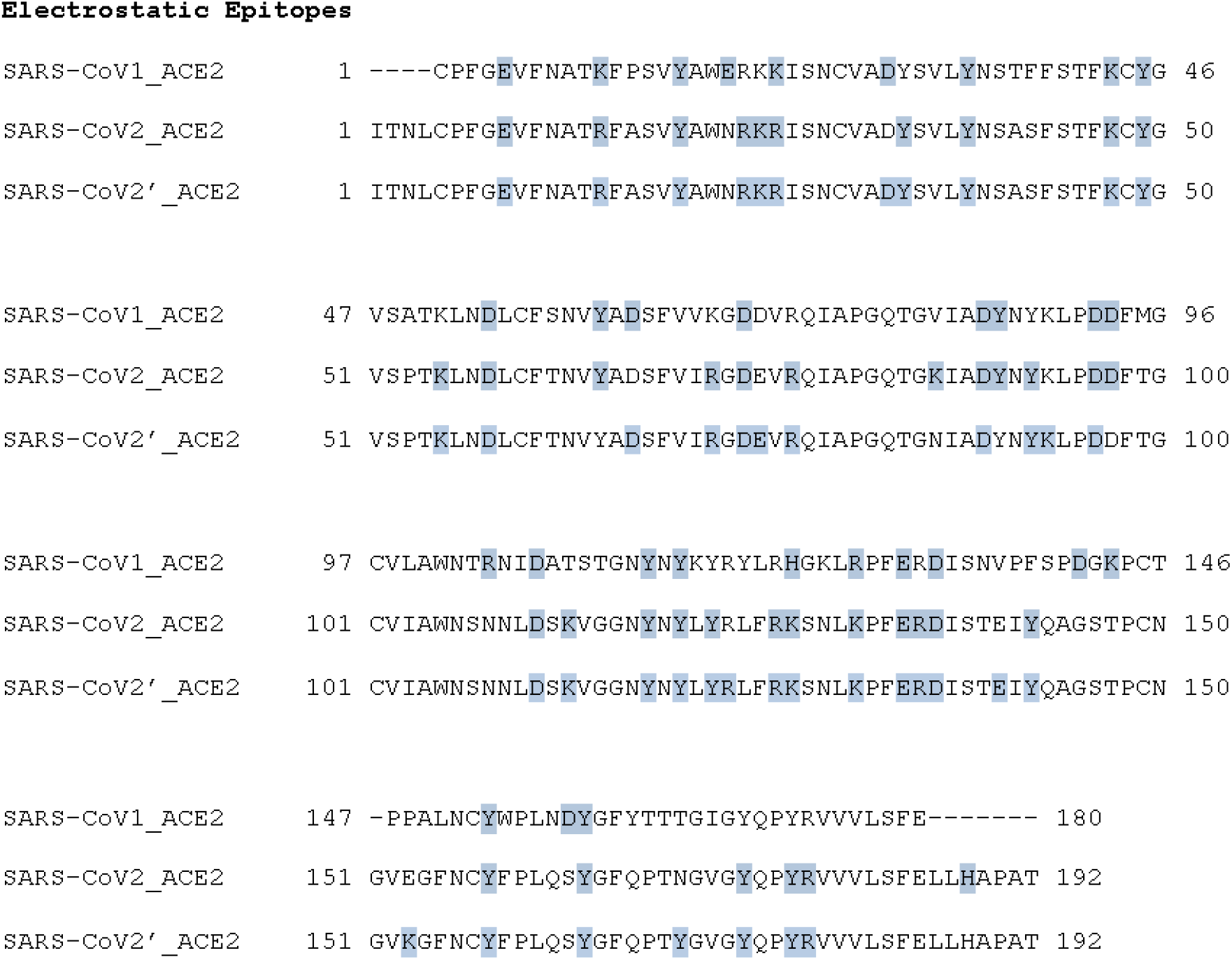
Electrostatic epitopes. Primary sequences of the SARS-CoV-1 S RBD, the SARS-CoV-2 S wt RBD, and the SARS-CoV-2’ – representing the South African variant (B.1.351) – with the interface with ACE2 (shown in blue). Data obtained using the threshold |Δp*Ka*|>0.01. Calculations for the RBD of SARS-CoV-2’ were performed with the structure RBD2_SA_. Gaps are represented by “-”. The numbers next to the chains are used to guide the identification of the amino acid sequence numbers of the RBD.

These results together with the amplified tendency for stronger complexation (see above) suggest that, as the virus SARS-CoV-2 evolved, the binding to the ACE2 receptor – which occurs with an increasing number of involved residues – became more specific. It is different from what is seen for the wildtype where the number of EE was larger than for SARS-CoV-1 but the affinity slightly weaker (a kind of “key-loose door lock cylinder” interaction). The rise in the specificity contributes to a higher virulence. This behaviour might also have an effect on how antibodies should block the RBD. If an antibody does not cover all the area given these EE, there is a chance that the RBD can still come closer to the ACE2 and probably allow the next steps of the infection to continue. This may reduce vaccine effectiveness.

### 3.2. The “up” state as a requirement for efficient binding

The spike protein hiddes several “tricks” via the changes on its conformational states. Wrapp *et al.*. (2020) revealed the movement of the RBD between the up/open and down/closed conformational states for SARS-CoV-2. This work was followed by many others that used experimental techniques to provide rich structural information regarding the interplay of the conformational transitions of the spike homotrimer (48,117,121). At least one chain of the homotrimer has to be at the “up” conformational state to allow the binding of the RBD with the receptor ACE2.

From a structural point of view, it is clear that most of the EE involved in the RBD-ACE2 complexation should be more exposed when the homotrimer is at the “up” conformational state. It is not only the steric clashes that are important to be removed for their binding. The key amino acids should be available for an effective interaction between these molecules. For this reason, we analysed the predicted EE for the RBD out of the trimer, and mapped them on the three dimensional macromolecular structures of the trimers. As can be seen in Table 2 for the SARS-CoV-2 wt RBD, the “down” conformational state (DDD) reduces the number of exposed (electrostatic) epitopes (NEE). The comparison of two experimental structures at different conformational states (DDD and UDD) indicates that there are 32 EE when one chain is at the “up” conformation (UDD state) while 25 are seen at the other conformational state. Interestingly, there is an increase in the NEE for the chains at the “down” state when at least one is at the “up” conformational state. This analysis was done using the detailed data for the mapping of EE for the RBD at different conformational states of the spike homotrimer at the amino acid level given in Table S.2.

**Table 2:**
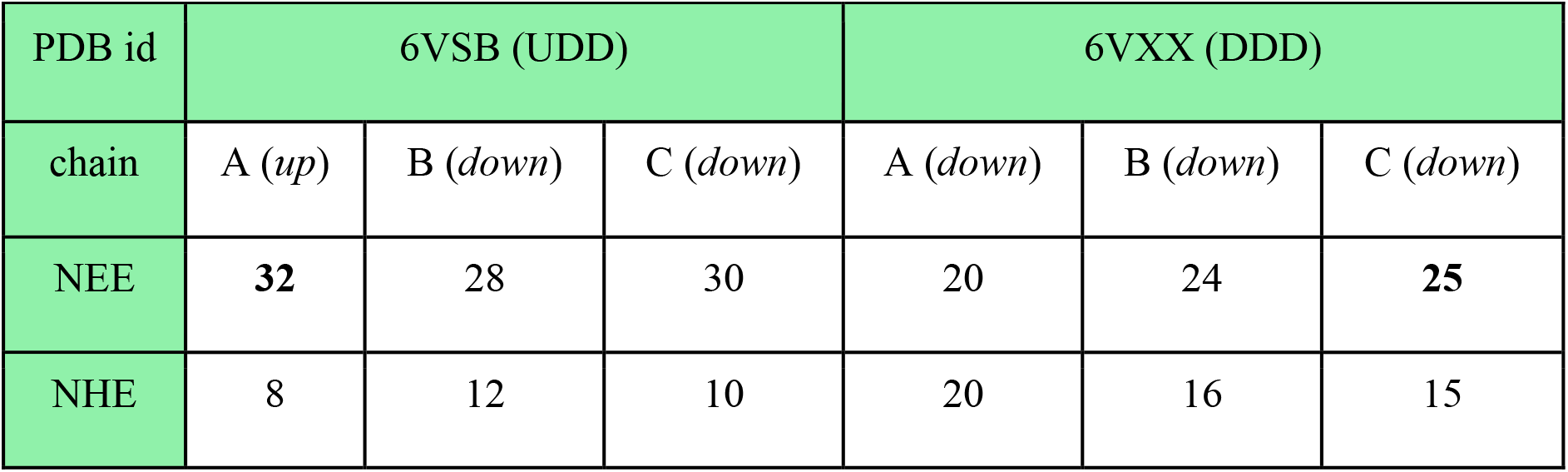
Location of the epitopes at different interfaces for SARS-CoV-2 wt S homotrimer. Number of exposed epitopes (NEE) and hidden epitopes (NHE) for interactions with ACE2 for two S homotrimer conformational states, one closed (DDD) and one open (UDD). This analysis used the PDB ids 6VXX and 6VSB, respectively. The state of each protein chain (D or U) is indicated in parentheses, where D corresponds to the closed (or “down”) conformational state, and U, to the open (or “up”) conformational state. The highest values are highlighted by bold fonts.

### 3.3. Electrostatic stability

Another molecular factor that can influence the transmissivity and virulence of SARS-CoV-2 through phenotypic changes is the increased stability of the spike homotrimers in the open state, exposing the EE of the RBD and, consequently, allowing the interaction between the RBD and ACE2 (41,43,54–56,58,59). Viruses often suffer mutations that improve either their binding affinities (as discussed above) or their proteins stabilities. An improvement on the affinities can result in a reduction in the stability which drives the needs of multiple mutations to keep or even increase the virulence. In principle, pH can have an important influence on this process too.

Therefore, we investigated (with the set 2 of the simulations) the electrostatic stability of the S trimeric structure of SARS-CoV-1, 2 wt and the SA variant as a function of pH for three different conformational states (namely, DDD, DUU and UDD). The aim was to identify which state has the greatest impact (favoring the upstream conformation of the Spike glycoprotein), and thus providing key information for the development of broader spectrum coping strategies against COVID-19.

Different analysis can be made exploring the stability of each protein sequence at different conformational states and the comparison between them. Starting with the sequence from SARS-CoV-1, its stability is greater (the values are more negative) over almost the entire extent of the relevant pH regimes when compared with SARS-CoV-2 wt as seen in Figure 4. The maximum estimated standard deviations on ΔG_elec_ for DDD, DUU and UDD are, respectively, 7, 14 and 10 kJ/mol for all studied pH. They were calculated based on the ten CpH simulations carried out with different coordinates of the trimer at the same conformational state. This lets the calculations to include some of the thermal fluctuations of the homotrimer structures. From pH 4.3 to 9.8, ΔG_elec_ is always negative for these three conformational states. Apparently, the most stable conformational state for this sequence (SARS-CoV-1) is DUU followed by UDD and DDD. Nevertheless, considering the differences between them with the estimated standard deviations, it does not allow us to identify the most stable one. For instance, at pH 7, ΔG_elec_ is equal to −61(4), −69(8) and −66(9) kJ/mol, respectively, for DDD, DUU and UDD. It was somehow unexpected to find that all the three conformational states would have statistically equivalent probabilities. Yet, there are twice more chances for the virus to be at the open form due to the fact that two states (DUU and UDD) offer this possibility. It infers that SARS-CoV-1 has a tendency to more often be ready to enter the human cell with at least one chain at the open state when its RBD is available to bind to the receptor ACE2 with great affinity (see discussion above). Also, it is tempting to make a connection between this molecular behavior with clinical observations. A higher probability to have the homotrimer at the “up” position should result in more symptomatic patients.

**Figure 4.**
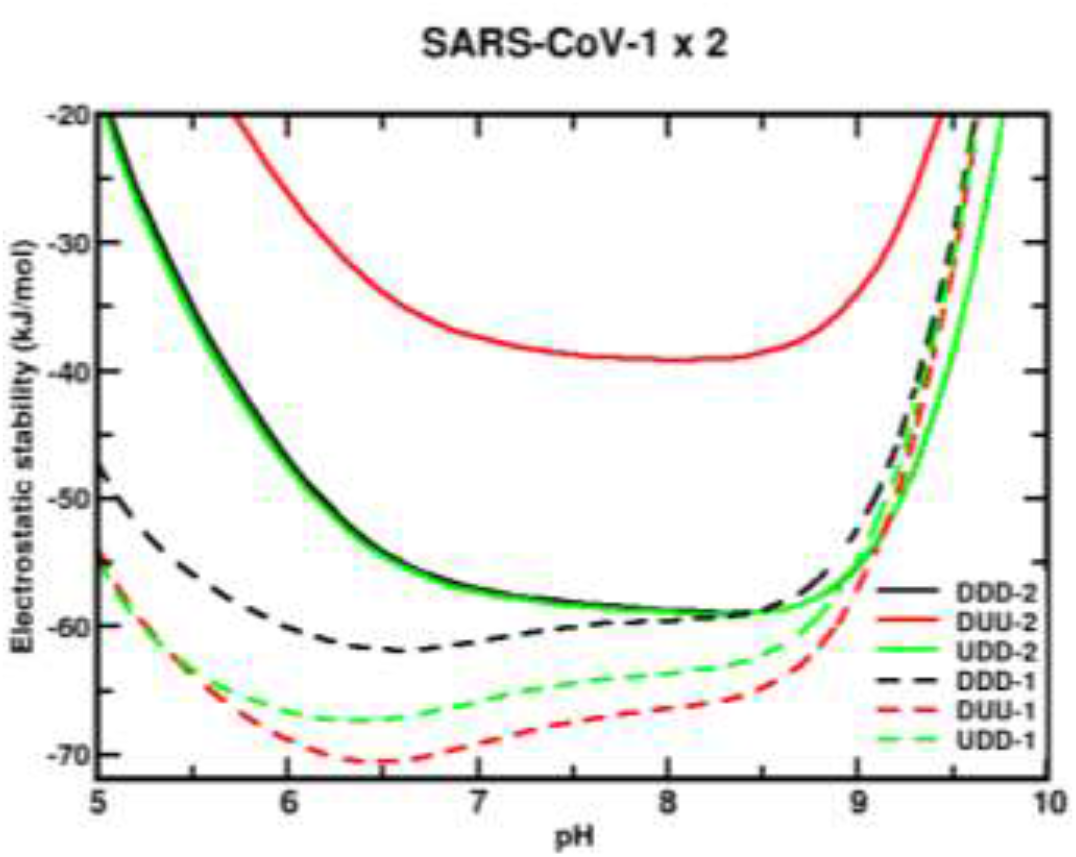
Simulated electrostatic stability profiles for the sequences of SARS-CoV-1 and SARS-CoV-2 wt spike homotrimers as a function of pH. The chains are in three different conformational states: DDD, DUU, and UDD, which are represented in black, red, and green, respectively. The continuous lines refer to SARS-CoV-1, and the dashed lines refer to SARS-CoV-2 wt. The three dimensional coordinates of the used structures in these CpH MC simulations were obtained from MD trajectories as described in the text. Each curve in this plot is an average of over 10 CpH simulations carried out with a trimer coordinate extracted from a different MD replica. Salt concentration was fixed at 150 mM.

pH has virtually a minor effect on the stability of this viral sequence for the most important biological regimes (pH ~4-7). The pH values where these states for SARS-CoV-1 are more stable are 6.6 (−62 kJ/mol), 6.5 (−71 kJ/mol), and 6.4 (−67 kJ/mol), respectively, for DDD, DUU and UDD. From pH 4 to 7, the differences in ΔG_elec_ [ΔΔG_elec_=ΔΔG_elec_(pH 7)- ΔΔG_elec_(pH 4)] for them is −103, −102 and −102 kJ/mol. The transitions from the close state (DDD) to the open states (UDD and DUU) require around 9 kJ/mol (depending on the final configurational state) at pH 4 whose value has the same order of magnitude as the estimated error. Another factor can help to trigger the “chameleon” behavior of the homotrimer. If experimentally solved CryoEM structures for the sequence given by SARS-CoV-1 [PDB ids 6ACC (DDD) and 6ACD (DUU)] are used in the calculations, the difference is relatively larger (19 kJ/mol) favouring the DDD state, although the unique structures for each state do not permit to estimate the errors as done for the MD coordinates (ΔG_elec_ = −100 kJ/mol, for DDD, and ΔG_elec_ = −71 kJ/mol, for DUU).

Conversely, all conformations of the sequence given by the SARS-CoV-2 wt are most stable at a solution pH between 6.5 and 9, being the most unstable conformation the one with two RBDs in the “up” position (i.e. DUU). The transition from DDD to DUU requires 20 kJ/mol at pH 7. Less is needed at lower pH regimes. The highest stability is observed for pHs 8.3 (−59 kJ/mol), 8.1 (−39 kJ/mol) and 8.3 (−59 kJ/mol) for, respectively, DDD, DUU and UDD. Furthermore, it can be seen that ΔG_elec_ for the whole homotrimer at the closed state (DDD) and with only one RBD (UDD) “up” is almost identical to each other (ΔG_elec(DDD)_ = −57±6 kJ/mol and ΔG_elec(UDD)_ = −57±9 kJ/mol at pH 7), showing a tendency for this state transition to be done without any energetic cost. Experimentally, the UDD was the most seen ACE2-bound conformational state (48). It can be hypothesised that this is the molecular reason that facilitates infections. An equal probability for DDD and UDD states as given by this theoretical analysis contributes to reduce the number of interactions RBD-ACE2 in comparison to what could be seen for SARS-CoV-1 (~ 66% for the two open states). Again, we can extrapolate our data suggesting that this can explain the molecular reasons for such a higher number of transmission by asymptomatic people as observed for SARS-CoV-2 and longer interval of contagiousness. In truth, the CryoEM study performed by Benton and others showed that 11% of the total trimeric structures were at the closed state (DDD) and 20% at the open state either with one RBD up (16%) or two (4%) (48). Also, most of the bound ACE2 cases in their work were observed for the UDD state (49% versus 14% for DUU and 3% for UUU).

Moreira and co-workers reported 10.4 kcal/mol and 32.5 kcal/mol for DDD→UDD and DDD→DUU transitions, respectively, at pH 7 and 150mM, using a Poisson-Boltzmann solver (84). No standard deviations were given. Our data agrees qualitatively only with the DDD→DUU transition. What triggers the conformational changes from one state to another (controlling the viral load in the patient) is not clear from these results. pH is not directly responsible for this transition, but it still can have an indirect contribution changing the interactions with another co-factor (e.g. glycans). This trigger mechanism was not addressed by our present simulations. It could be that pH promotes intermediate conformational states which can only be properly described by a full CpH MD simulations whose computational costs are still prohibitive.

We next analysed the sequence effects of the SA variant (B.1.351) comparing it with the wildtype of SARS-CoV-2. This comparison revealed a significant improvement in the hometrimer stability for the SA variant, although the stability curves are still qualitatively similar (see Figure 5). Unlike the wildtype, the SA variant presents the closed (DDD) state as the less favorable one, while the UDD and DUU forms have similar values. However, the most relevant information is the high difference of approximately 175 kJ/mol between the most stable curve of the wildtype and this variant. This shows how much more stable the SA variant (B.1.351) is in conformational states that enables it to easily infect the human cells. The mutations D80A, D215G and R246I are probably the main responsible for this higher stability. In fact, these substitutions contribute to make the trimer more positively charged. Comparing the net charges of the wildtype with the SA variant, we find that they increased from +4.6 to +10.6, for DDD, from +4.7 to +10.7, for DUU, from +4.6 to +10.6, for UDD, at pH 7.

**Figure 5.**
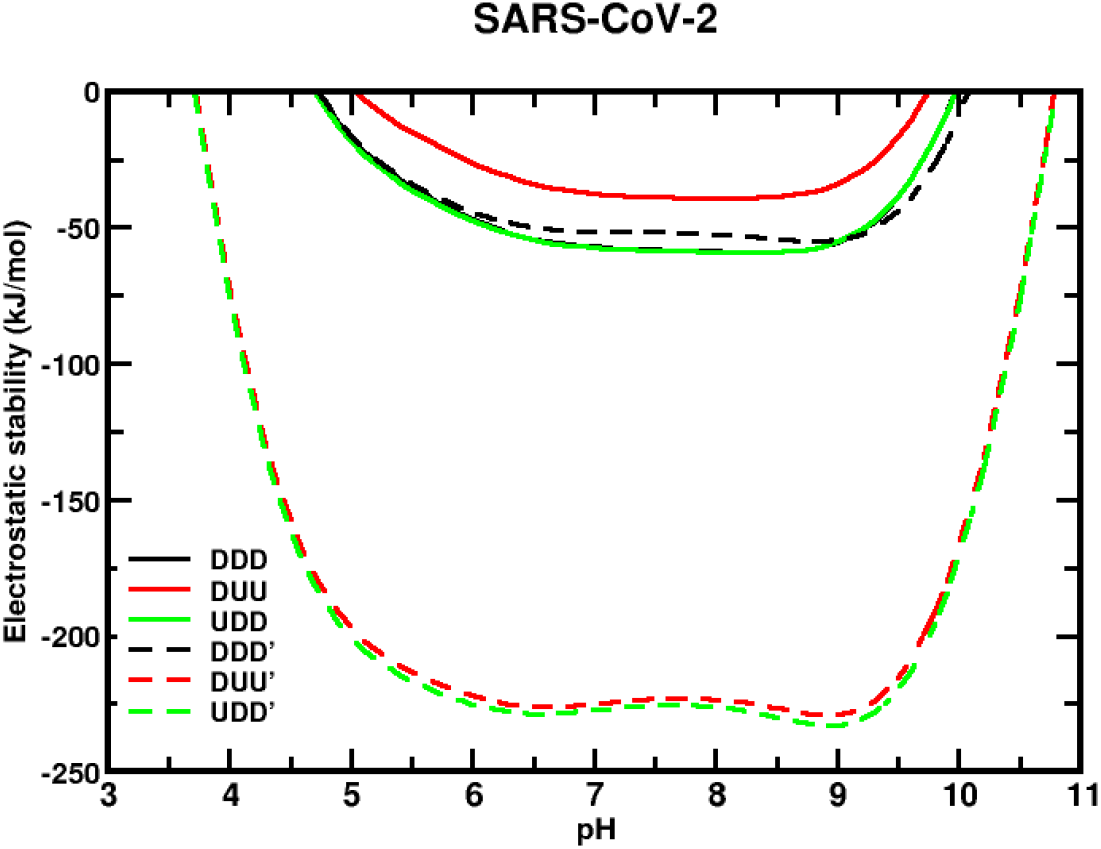
Simulated electrostatic stability profiles for the sequences of the SARS-CoV-2 wt and the South African (B. 1.351) spike homotrimers as a function of pH. The chains are in three different conformational states: DDD, DUU, and UDD, which are represented in black, red, and green, respectively. The continuous lines refer to SARS-CoV-2 wt, and the dashed lines refer to South African variant (B.1.351). All other details as in Figure 4.

Based on this data, it appears that evolution has switched off its “chameleon” feature dismissing the need for the DDD state due to the facility for people being contaminated. The SA variant seems to be always ready to let the RBD be prepared to form a complex with ACE2 explaining how much more harmful to society it can be. If the hypothesis that an equal probability for DDD and UDD implies more asymptomatic cases for the wildtype (see Figure 4), the increased probability here for the two open states of the SA sequence should decrease the percentage of asymptomatic and increase the number of symptomatic ones. As the time passes, more epidemiological data might be available to corroborate this hypothesis.

The stability of the trimer with one ACE2-bound was also investigated to assess the contribution that the receptor ACE2 could have on it. Bai and Warshel noticed that the RBD of SARS-CoV-2 has been optimized to bind stronger at distant sites (122). They also questioned whether this stronger binding could be related with conformational changes of the homotrimer. This can be partially answered by this analysis. Here, an experimental CryoEM structure (PDB id 7A94) was used for these calculations. This structure corresponds to the UDD conformational state. In Figure 6, it is shown ΔG_elec_ for the sequence given by SARS-CoV-2 wildtype complexed with one ACE2 molecule (holo state). In the same Figure 6, ΔG_elec_ for the homotrimer at the *same* conformation but with the ACE2 molecule removed (apo state) is plotted. The trimer conformation at the apo state is a bit more elongated than the ones used in the previous calculations discussed above with the structures from MD. The RMSD between this trimer structure and one from the MD trajectories is 1.3Å. This explains why ΔGele dropped to values much smaller than what was seen above. Indeed, this suggests that approaching the receptor ACE2 modifies the trimer conformation and turns its structure to a more stable one. Again, another molecular mechanism to promote the infection. Although not explicitly investigated here, this could also be one trigger to promote the conformational changes from the DDD to UDD (or DUU). By comparing ΔG_elec_ for the homotrimer at the halo and apo states, the effect of ACE2 in the stability is finally elucidated. As expected, ACE2 increases further the stability of the trimer, favouring even more the complexation and the cell entry itself as can be seen in Figure 6. These structures are more stable around pH 6.3 [pH 6.4 (ΔG_elec_=-254 kJ/mol) for the trimer and pH 6.2 (ΔG_elec_=-323 kJ/mol) for the complex]. The ACE2-bound form of the trimer favours its stability. At pH 7, the difference between the holo and apo forms is −59 kJ/mol. This value might be underestimated because we are comparing the trimer in an already modified conformation by the previous presence of ACE2. If the initial state is the UDD obtained in the absence of ACE2 (as the ones generated at the MD trajectory), the difference is increased to −249 kJ/mol. Although the physical chemical conditions of the two trimer structures (the CryoEM and the MD) are not the same, this difference should reflect at least the order of magnitude of the energetic gain when the trimer goes from a single molecule to a complex with ACE2.

**Figure 6.**
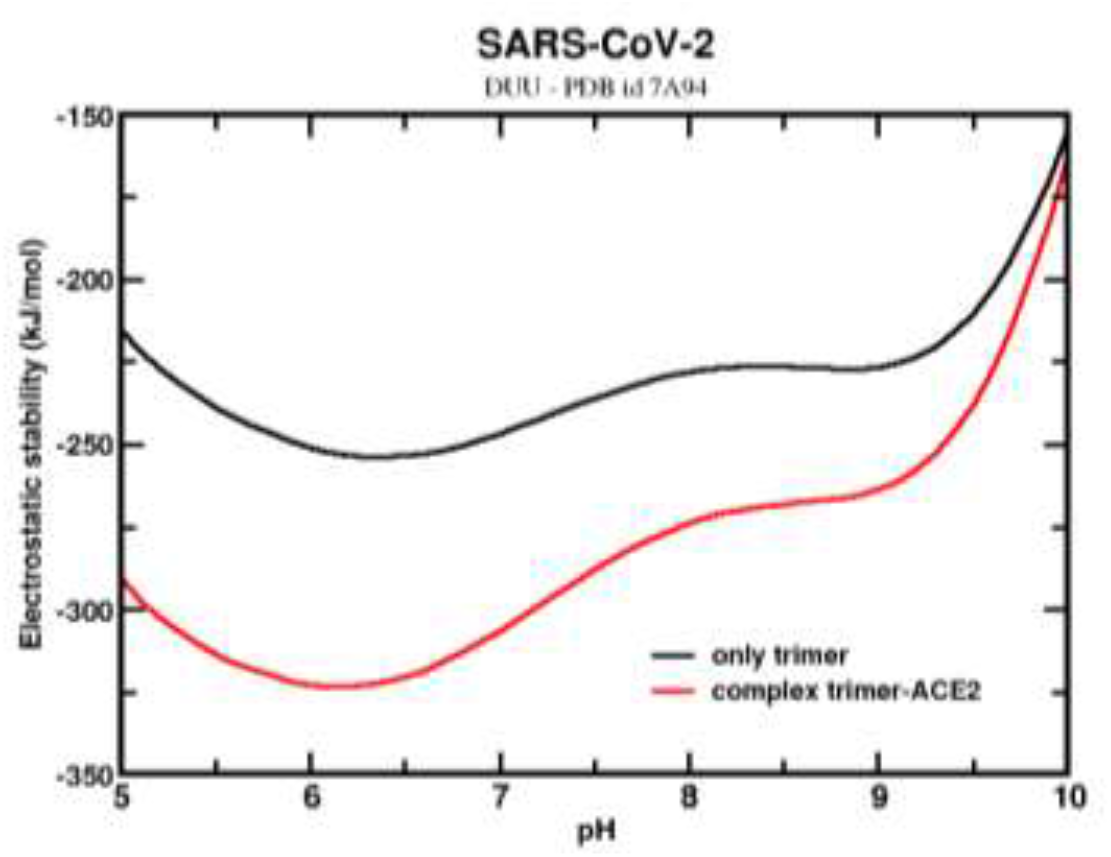
Simulated electrostatic stability profiles for the SARS-CoV-2 wt spike homotrimers at two different ACE2-bound forms as a function of pH. The black continuous line refers to SARS-CoV-2 wt at the apo state (maintaining the same conformation of the complex), and the red line refers to the complex trimer-ACE2. The three-dimensional coordinate was given by the PDB id 7A94. Salt concentration was fixed at 150 mM.

## 4. Discussion

Some molecular aspects relevant for the understanding of the increased transmissivity and virulence of new variants of SARS-CoV-2 were discussed. This study involved complementary pieces of information: 1.) the theoretical estimation of the binding affinities between the RBD of the Spike proteins from different mutants with the cellular receptor ACE2, 2.) the mapping of the electrostatic epitopes, and 3.) the electrostatic stability of different conformational states of the whole spike homotrimer. In particular, we identified for the studied strains the probability to have the trimer at the open state, since the interaction with ACE2 only occurs if the epitopes are available for binding. Even though these mechanisms may be connected by unknown factors, we could identify some key aspects of the natural evolution of this virus.

All studied variants showed a tendency for a higher RBD-ACE2 binding affinities when compared with the wild type version of SARS-CoV-2. Once the analysis of the complexion between ACE2 and the RBD from the SA variant (B.1.351) showed minimal increases in the free energy of interactions (despite an increase in the specificity with more amino acids that are important for this process), we investigated another aspect that is often used by viruses upon mutations: the stability. Comparing three conformational states (DDD, UDD and DUU) for SARS-CoV-1, SARS-CoV-2 wt and its SA variant, we could see differences on the homotrimer probabilities for being at these states. From SARS-CoV-1 to SARS-CoV-2 wt, there is a slightly smaller chance for the SARS-CoV-2 trimer to be available for binding with ACE2. The increase in the probability for DDD was also observed and interpreted as a possible explanation for a higher number of asymptomatic patients and longer interval of contagiousness. Conversely, the SA variant does favor the open state of the trimer. At pH 7, the stability of the trimer is ca. 6 times higher for the SA variant (ΔG_elec[wt]_=-72±4 kJ/mol and ΔG_elec[SA]_=-225±4 kJ/mol for DUU, and ΔG_elec[wt]_=-75±5 kJ/mol and ΔG_elec[SA]_=-227±5 kJ/mol for UDD). The synergistic effect between the tendency to increase the affinity with ACE2 and the availability of RBD up for binding promotes higher transmissivity and virulence for the SA variant. This might be even more substantial that we could estimate here due all the approximations assumed. For instance, we did not touch on other effects that can be affected by pH and amplify more the virulence [e.g. the contributions from the glycams; the interaction of the viral proteins with other cellular receptors, such as TMPRSS2, which cleaves protein S at two sites, and NEUROPILIN-1, an alternative host factor for SARS-CoV-2 (123,124)].

Viruses are simply a group of organisms that exploits their environment in order to survive, thus they have to find a balance between the harm it causes and its ability to transmit itself (125). By looking historically at the human diseases caused by the coronavidae family, the Severe Acute Respiratory Disease from 2003, caused by SARS-CoV-1, did not cause such disastrous effects because, even though the case fatality rate was 9.6%, the disease caused such deleterious effects on the body that the infected individuals isolated themselves in their homes. (14) This might be interpreted by the disponibility of spike homotrimers for being at the open state (DUU and UDD) as we discussed above. Transmissivity was also affected because the virus was not able to transmit itself to a large number of people.

SARS-CoV-2, on the other hand, developed a mechanism that enabled him to stay in a closed state in order to silently spread more widely – a kind of “chameleon” feature as cited above. As a result, the number of cases raised alarmingly, and the average of the COVID-19 case fatality rate is about 2%–3% worldwide (126). As time went by and the majority of the population stopped following the security measures, the VOCs included mutations that dismissed the practical need of the “chameleon” feature (probably due to a higher probability of trimers at the DDD conformation state), made them more virulent and dangerous. The combination of all the mechanisms listed above gave the SA variant a well improved fitness from the virus perspective. Even though there are still very few studies on the matter, the case fatality rate is expected to increase.

Besides shedding some light on how the virus uses physical chemical properties to evolve, the molecular mechanisms reported here have also direct implications for physiopathology, therapeutic strategies and vaccines. The amount of available receptors ACE2 can be easily compensated by an increased affinity and elevated number of receptors ready to dock at the human cell. Higher RBD-ACE2 binding affinities observed for the new variants imply that younger infected individuals without comorbidities and with naturally less disponibility of receptors ACE2 can have similar clinical conditions as observed for older patients contaminated by the wild type version.

The neutralization of the new variants would require an elevated number of binders. For instance, it has to be investigated if higher doses of the available vaccines could boost the immune system to produce the enough concentration of antibodies to neutralize such a high number of RBDs ready to interact with ACE2. Considering the increased stability of the B. 1.351variant in its infectious form, the need to adopt prompt and effective measures to contain the advance of the pandemic must be reinforced. Even though one is supposed to have protection in some level with vaccination against the early variants, the necessary concentration of antibodies for the new variants might not be achieved so fast by the human body. Social distance, masks, and rapid and global mass vaccination should be adopted to prevent transmission of the virus, since it only tends to accumulate mutations to be more transmittable and more prepared to evade the body’s immune strategies (39). It is worth noting that the strongest ACE2-affinity-enhancing mutations have not been yet selected in current variants. As the virus evolves it can still find other substitutions to improve its destructive power to spread and thrive (from his perspective) increasing the risk to our survival. (46).

## Author Contributions

C. C. G.: Visualization, Investigation, Writing – original draft. F. L. B. da S.: Conceptualization, Investigation, Methodology, Software, Data curation, Writing – review & editing, Supervision. A. L.: Data curation, Methodology, Writing – review & editing, Supervision.

## Acknowledgments

This work has been supported by the “Fundação de Amparo à Pesquisa do Estado de São Paulo” [Fapesp 2020/07158-2 (F.L.B.d.S.)] and the “Conselho Nacional de Desenvolvimento Científico e Tecnológico” [CNPq 305393/2020-0 (F.L.B.d.S.) and PIBIC/CNPq 2020-1732 (C.C.G.)]. F.L.B.d.S. is also thankful to the resources provided by the Swedish National Infrastructure for Computing (SNIC) at NSC. It is also a pleasure to acknowledge initial discussions with Adolfo Pomo (researcher at the Institute of Fundamental Technological Research PAN, Poland). A.L. acknowledges Swedish Science Council for financial support, and partial support from a grant from the Ministry of Research and Innovation of Romania (CNCS – UEFISCDI, project number PN-III-P4-ID-PCCF-2016-0050, within PNCDI III).

