## SupplementaryMaterial for "The Up state of the SARS-COV-2 Spike homotrimer favors an increased virulence for new variants"

**Supplementary Material**

Additional details are given as supplementary material for:

1. **Supplementary Figures**

Sequences alignment of SARS-CoV-1 and SARS-CoV-2 RBD (S1 subunit) proteins

```
SARS-CoV-1         1 ----CPFGEVFNATKFPSVYAWERKKISNCVADYSVLYNSTFFSTFKCYG     46
                         ||||||||||:|.|||||.||:||||||||||||||..||||||||
SARS-CoV-2         1 ITNLCPFGEVFNATRFASVYAWNRKRISNCVADYSVLYNSASFSTFKCYG     50

SARS-CoV-1        47 VSATKLNDLCFSNVYADSFVVKGDDVRQIAPGQTGVIADYNYKLPDDFMG     96
                     ||.||||||||:||||||||::||:||||||||||.||||||||||||.|
SARS-CoV-2        51 VSPTKLNDLCFTNVYADSFVIRGDEVRQIAPGQTGKIADYNYKLPDDFTG    100

SARS-CoV-1        97 CVLAWNTRNIDATSTGNYNYKYRYLRHGKLRPFERDISNVPFSPDGKPCT    146
                     ||:|||:.|:|:...|||||.||..|...|:|||||||...:.....||.
SARS-CoV-2       101 CVIAWNSNNLDSKVGGNYNYLYRLFRKSNLKPFERDISTEIYQAGSTPCN    150

SARS-CoV-1       147 -PPALNCYWPLNDYGFYTTTGIGYQPYRVVVLSFE-------    180
                      ....|||:||..|||..|.|:|||||||||||||
SARS-CoV-2       151 GVEGFNCYFPLQSYGFQPTNGVGYQPYRVVVLSFELLHAPAT    192
```

**Figure S.1.** Sequence alignment of SARS-CoV-1 and SARS-CoV-2 wt RBD (S1 subunit) proteins. The numbers next to the chains are used to guide the identification of the amino acid sequence numbers of the RBD. All sequence numbers were automatically shifted by the EMBOSS Needle server. Original numbers can be recovered by the addition of 331 for the SARS-CoV-2 (for instance, N170 in this alignment corresponds to N501 in the PDB files for this strain). The signs between the aligned sequences have the typical meaning: a) "|" indicates equal amino acids in both sequences; b) ";" indicates similarities with a high score and c) "." indicates the amino acids with a low positive score. Gaps are represented by "-". See text for more information.

Sequences alignment of SARS-CoV-2 and SARS-CoV-2' RBD (S1 subunit) proteins

```
SARS-CoV-2 wt              1 ITNLCPFGEVFNATRFASVYAWNRKRISNCVADYSVLYNSASFSTFKCYG     50
                             ||||||||||||||||||||||||||||||||||||||||||||||||||
SARS-CoV-2 (B.1.351)       1 ITNLCPFGEVFNATRFASVYAWNRKRISNCVADYSVLYNSASFSTFKCYG     50

SARS-CoV-2 wt             51 VSPTKLNDLCFTNVYADSFVIRGDEVRQIAPGQTGKIADYNYKLPDDFTG    100
                             |||||||||||||||||||||||||||||||||||.||||||||||||||
SARS-CoV-2 (B.1.351)      51 VSPTKLNDLCFTNVYADSFVIRGDEVRQIAPGQTGNIADYNYKLPDDFTG    100

SARS-CoV-2 wt            101 CVIAWNSNNLDSKVGGNYNYLYRLFRKSNLKPFERDISTEIYQAGSTPCN    150
                             ||||||||||||||||||||||||||||||||||||||||||||||||||
SARS-CoV-2 (B.1.351)     101 CVIAWNSNNLDSKVGGNYNYLYRLFRKSNLKPFERDISTEIYQAGSTPCN    150

SARS-CoV-2 wt            151 GVEGFNCYFPLQSYGFQPTNGVGYQPYRVVVLSFELLHAPAT    192
                             ||:||||||||||||||||.||||||||||||||||||||||
SARS-CoV-2 (B.1.351)     151 GVKGFNCYFPLQSYGFQPTYGVGYQPYRVVVLSFELLHAPAT    192
```

**Figure S.2.** Sequence alignment of SARS-CoV-2 wt and SARS-CoV-2 (B.1.351) RBD (S1 subunit) proteins. All the other details are as in Fig. S1.

Sequences alignment of SARS-CoV-1 and SARS-CoV-2’ RBD (S1 subunit) proteins

```
SARS-CoV-1                  1 ----CPFGEVFNATKFPSVYAWERKKISNCVADYSVLYNSTFFSTFKCYG     46
                                  ||||||||||:|.|||||.||:||||||||||||||..||||||||
SARS-CoV-2 (B.1.351)        1 ITNLCPFGEVFNATRFASVYAWNRKRISNCVADYSVLYNSASFSTFKCYG     50

SARS-CoV-1                 47 VSATKLNDLCFSNVYADSFVVKGDDVRQIAPGQTGVIADYNYKLPDDFMG     96
                              ||.||||||||:||||||||::||:||||||||||.||||||||||||.|
SARS-CoV-2 (B.1.351)       51 VSPTKLNDLCFTNVYADSFVIRGDEVRQIAPGQTGNIADYNYKLPDDFTG    100

SARS-CoV-1                 97 CVLAWNTRNIDATSTGNYNYKYRYLRHGKLRPFERDISNVPFSPDGKPCT    146
                              ||:|||:.|:|:...|||||.||..|...|:|||||||...:.....||.
SARS-CoV-2 (B.1.351)      101 CVIAWNSNNLDSKVGGNYNYLYRLFRKSNLKPFERDISTEIYQAGSTPCN    150

SARS-CoV-1                147 -PPALNCYWPLNDYGFYTTTGIGYQPYRVVVLSFE-------    180
                               ....|||:||..|||..|.|:|||||||||||||
SARS-CoV-2 (B.1.351)      151 GVKGFNCYFPLQSYGFQPTYGVGYQPYRVVVLSFELLHAPAT    192
```

**Figure S.3.** Sequence alignment of SARS-CoV-1 and SARS-CoV-2 (B.1.351) RBD (S1 subunit) proteins. All the other details are as in Fig. S1.

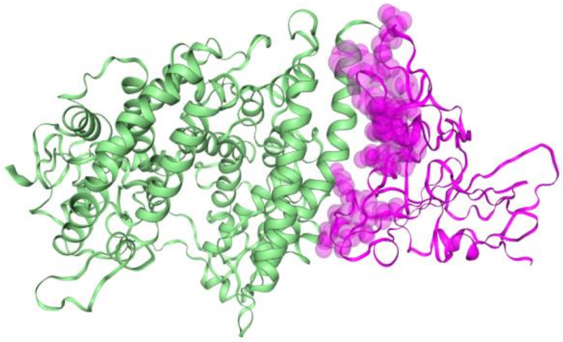

**Figure. S.4.** Molecular structures of the SARS-CoV-2 S RBD complexed with ACE2. For better visualization of the epitope-paratope interface, the interface residues on the RBD were represented as overpost pink spheres. Interface residues are defined as residues whose atoms are within a distance of 5 Angstroms from other atoms of the neighboring chains. Magenta represents RBD on SARS-CoV-2 and the light green represents the receptor ACE2 (PDB id 6LZG). The image was generated by CoV3D (Gowthaman *et al.*, 2021).

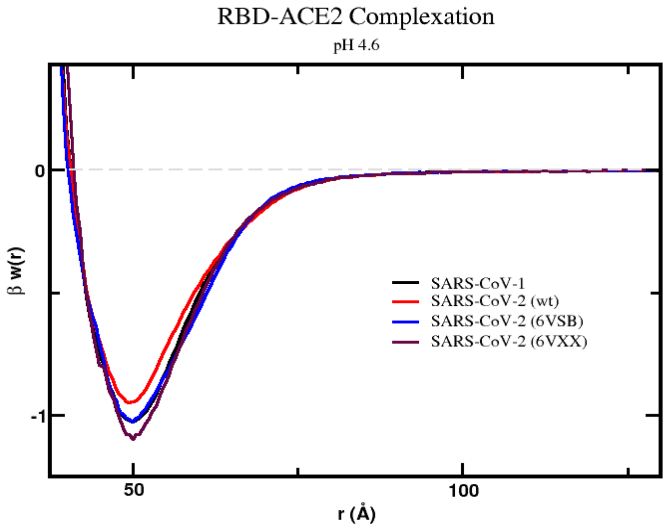


**Figure. S.5**. Free energy profiles for the interaction of RBD proteins obtained from different structural data with the cellular receptor ACE2. The simulated free energy of interactions [$\boldsymbol{\beta} w(r)$] between the centers of the RBD proteins from SARS-CoV-1 and different coordinates for the SARS-CoV-2 wildtype interacting with the cellular receptor ACE2 are given at pH 4.6. The source of the three dimensional structures of these proteins are explained in the text and referred as $RBD1_{wt}$, $RBD2_{wt,}$, $RBD2_{wt'(6VSB),}$and $RBD2_{wt'(6VXX)}$, respectively. The plots for $RBD1_{wt}$ and $RBD2_{wt,}$ correspond to the same ones already given in Figure 2. Salt concentration was fixed at 150 mM. Simulations started with the two molecules placed at random orientation and separation distance. Results for SARS-CoV-1, SARS-CoV-2 (wt), SARS-CoV-2 (wt with RBD from PDB id 6SVB), and SARS-CoV-2 (wt with RBD from PDB id 6SXX) are shown as black, red, dark blue, and marron continuous lines, respectively.

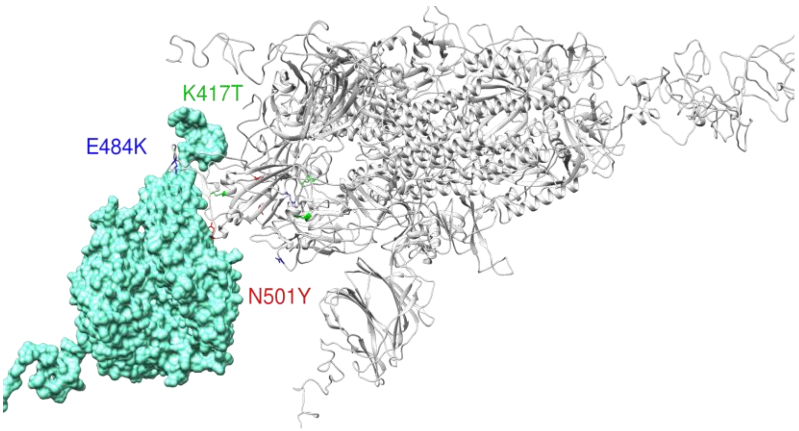

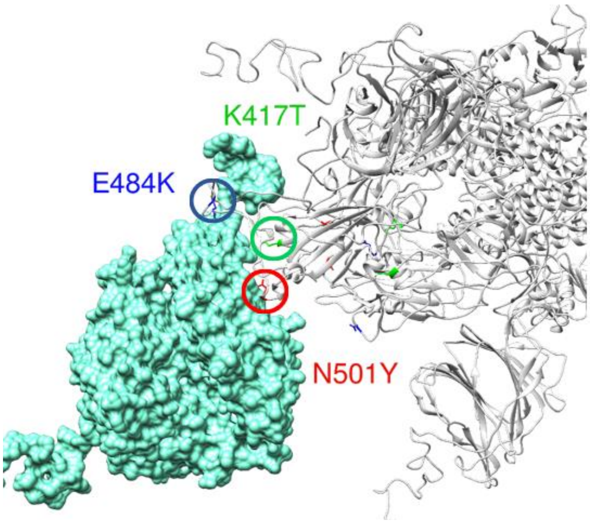


**Figure S.6.** Molecular structures of the SARS-CoV-2 Spike complexed with ACE2. The aquamarine surface represents the receptor ACE2 and the grey ribbon represents the Spike homotrimer protein (PDB id 7A94). The top image shows the full homotrimer with ACE2 while the zoom is given to the RBD region at the bottom one. For better visualization of the epitope-paratope interface, the Spike residue N501 is shown in red, E484 in blue, and K417 in green. The amino acids are substituted by Y, K and T, respectively, for the Brazilian P.1 variant. The image was generated by UCSF Chimera (Pettersen *et al.*, 2004).

### 2. Supplementary Tables

**Table S.1**. Main physical chemical properties of the RBDs of different new variants. These results were obtained by the FPTS at physiological salt concentration. See the text for more details.

| | Charges numbers for different protein structures | | | | | | | |
|---|---|---|---|---|---|---|---|---|
| | RBD2$_{wt}$ | RBD2$_{m}$ | RBD2$_{SA}$ | RBD2$_{BR}$ | RBD2$_{CA}$ | RBD2$_{NY}$ | RBD2$_{UK}$ | RBD2$_{I}$ |
| pH 4.6 | 5.5 | 5.5 | 6.2 | 6.2 | 6.4 | 7.1 | 5.5 | 7.0 |
| pH 7.0 | 2.2 | 2.2 | 3.2 | 3.2 | 3.1 | 4.1 | 2.1 | 4.1 |
| | Dipole moment numbers for the same structures | | | | | | | |
| pH 4.6 | 35 | 35 | 65 | 65 | 40 | 71 | 35 | 53 |
| pH 7.0 | 31 | 31 | 82 | 82 | 43 | 90 | 31 | 71 |

**Table S.2.** Detailed data for the mapping of EE for the RBD at different conformational states of the spike homotrimer. The EE were predicted by the PROCEEDpKa method for the RBD of SARS-CoV-2 wt out of the homotrimer (as given in Figure 3). Exposed and hidden residues were identified by the on-line server "PDBePISa" (90) with default options. PDB ids 6VXX and 6VSB were used in this analysis.

a) EEs exposed to the solvent and available to directly interact with the receptor ACE2 for chain A of Spike RBD (PDB id 6VSB).

| Spike RBD EE - Chain A (NEE=32) (NHE=6) (*up*) | | | |
|---|---|---|---|
| E340 | K386 | D427 | E465 |
| R346 | D389 | D428 | R466 |
| Y351 | Y396 | D442 | D467 |
| R355 | R403 | K444 | Y473 |
| K356 | D405 | Y449 | Y489 |
| R357 | R408 | Y451 | Y495 |
| Y365 | K417 | Y453 | Y505 |
| Y369 | D420 | R457 | Y508 |
| K378 | Y421 | K458 | R509 |
| Y380 | Y423 | K462 | H519 |

b) EEs exposed to the solvent and available to directly interact with the receptor ACE2 for chain B of Spike RBD (PDB id 6VSB).

| Spike RBD EE - Chain B (NEE=28) (NHE=11) (*down*) | | | |
|---|---|---|---|
| E340 | K386 | D427 | E465 |
| R346 | D389 | D428 | R466 |
| Y351 | Y396 | D442 | D467 |
| R355 | R403 | K444 | Y473 |

| | | | |
|---|---|---|---|
| K356 | D405 | Y449 | Y489 |
| R357 | R408 | Y451 | Y495 |
| Y365 | K417 | Y453 | Y505 |
| Y369 | D420 | R457 | Y508 |
| K378 | Y421 | K458 | R509 |
| Y380 | Y423 | K462 | H519 |

c) EEs exposed to the solvent and available to directly interact with the receptor ACE2 for chain C of Spike RBD (PDB id 6VSB).

| Spike RBD EE - Chain C (NEE=30) (NHE=10) (*down*) | | | |
|---|---|---|---|
| E340 | K386 | D427 | E465 |
| R346 | D389 | D428 | R466 |
| Y351 | Y396 | D442 | D467 |
| R355 | R403 | K444 | Y473 |
| K356 | D405 | Y449 | Y489 |
| R357 | R408 | Y451 | Y495 |
| Y365 | K417 | Y453 | Y505 |
| Y369 | D420 | R457 | Y508 |
| K378 | Y421 | K458 | R509 |
| Y380 | Y423 | K462 | H519 |

d) EEs exposed to the solvent and available to directly interact with the receptor ACE2 for chain A of Spike RBD (PDB id 6VXX).

| Spike RBD EE - Chain A (NEE=20) (NHE=20) (*down*) | | | |
|---|---|---|---|
| E359 | K405 | D446 | E484 |

| | | | |
|---|---|---|---|
| R365 | D408 | D447 | R485 |
| Y370 | Y415 | D461 | D486 |
| R374 | R422 | K463 | Y492 |
| K375 | D424 | Y468 | Y508 |
| R376 | R427 | Y470 | Y514 |
| Y384 | K436 | Y472 | Y524 |
| Y388 | D439 | R476 | Y527 |
| K397 | Y440 | K477 | R528 |
| Y399 | Y442 | K481 | H538 |

e) EEs exposed to the solvent and available to directly interact with the receptor ACE2 for chain B of Spike RBD (PDB id 6VXX).

| Spike RBD EE - Chain B (NEE=24) (NHE=15) (*down*) | | | |
|---|---|---|---|
| E359 | K405 | D446 | E484 |
| R365 | D408 | D447 | R485 |
| Y370 | Y415 | D461 | D486 |
| R374 | R422 | K463 | Y492 |
| K375 | D424 | Y468 | Y508 |
| R376 | R427 | Y470 | Y514 |
| Y384 | K436 | Y472 | Y524 |
| Y388 | D439 | R476 | Y527 |
| K397 | Y440 | K477 | R528 |
| Y399 | Y442 | K481 | H538 |

f) EEs exposed to the solvent and available to directly interact with the receptor ACE2 for chain C of Spike RBD (PDB id 6VXX).

| Spike RBD EE - Chain C (NEE=25) (NHE=15) (*down*) | | | |
|---|---|---|---|
| E359 | K405 | D446 | E484 |
| R365 | D408 | D447 | R485 |
| Y370 | Y415 | D461 | D486 |
| R374 | R422 | K463 | Y492 |
| K375 | D424 | Y468 | Y508 |
| R376 | R427 | Y470 | Y514 |
| Y384 | K436 | Y472 | Y524 |
| Y388 | D439 | R476 | Y527 |
| K397 | Y440 | K477 | R528 |
| Y399 | Y442 | K481 | H538 |

**Legend**

| | |
|---|---|
| | Solvent-accessible residues |
| | Inaccessible residues |
| | Interfacing residues (AxB) |
| | Interfacing residues (AxC) |
| | Interfacing residues (BxC) |
| **NEE** | Number of exposed epitopes |
| **NHE** | Number of hidden epitopes (due to the interface with another chain) |
